# Genome-wide latitudinal variation associated with defence and reproductive pathways linked to Norway spruce adaptation

**DOI:** 10.64898/2026.11.06.764856

**Authors:** Sonali Sachin Ranade, María Rosario García-Gil

**Affiliations:** Umeå Plant Science Centre (UPSC), Department of Forest Genetics and Plant Physiology, Swedish University of Agricultural Sciences, 901 83 Umeå, Sweden

**Keywords:** Cline, cSNP, defence, flowering, local adaptation, natural variation, Norway spruce

## Abstract

Environmental cues modulate the natural variations in genes that outline local adaptation in plants for optimal survival. Genome-wide analysis using exome capture dataset in Norway spruce (*Picea abies* L. Kast) in Sweden revealed natural allelic variation in coding regions of genes associated with defence and reproductive pathways that follow a latitudinal cline, despite of the weak population structure with a low genetic differentiation. Steepest clines in the allele frequencies of missense SNPs from coding region (cSNPs) were detected in *60S RIBOSOMAL PROTEIN L18* (*RPL18C*), a gene involved in plant survival and defence/stress responses and *LONG AFTER FAR-RED 3* (*LAF3*) that mediates light responses. This was closely followed by cline in *ZINC 4 FINGER DNA 3’-PHOSPHOESTERASE* (*ZDP*), involved in DNA repair and DNA demethylation pathways. Flowering genes that displayed latitudinal cline in cSNPs included *BARELY ANY MERISTEM 1* (*BAM1*), *REDUCED VERNALIZATION RESPONSE 1* (*VRN1*), *FLOWERING CONTROL LOCUS A* (*FCA*) and *TERMINAL FLOWER 1* (*TFL1*) among few others. These variations may contribute to differential functioning of the defence/flowering genes, enabling the tree to thrive at its best under given local environmental conditions. The clines provide evidence towards local adaptation in these Norway spruce populations across Sweden; it coincides with the latitudinal variation in climatic conditions including temperature, day length and light quality, along with existing cline e.g. differential growth and defence response, and bud-burst/bud-set. The knowledge can be applied in forest industry to design future sustainable breeding strategies e.g. assisted migration, in the context of inevitable climate change.

**Highlights:** Genome-wide analysis from exome capture revealed natural variation in genes associated with defence and reproductive pathways in Norway spruce population across Sweden. Variation in allele frequencies of SNPs that follows a latitudinal cline, despite of weak population structure with low genetic differentiation, may represent cues for local adaptation. Knowledge gained can be applied to design future sustainable breeding strategies in forestry.

## Introduction

A latitudinal cline is defined as a measurable gradient of a single trait or character in a particular species, across its geographical range which is shaped by natural selection matching the existing climatic conditions. Clines provide information on patterns of genetic adaptation to varying environmental factors e.g. temperature, photoperiod and light quality. Local adaptation occurs when resident populations evolve divergent phenotypic traits allowing the organism to thrive optimally under the prevailing environmental conditions including biotic and abiotic factors. Genome-wide screening for latitudinal clines is an efficient method that can be applied to identify novel genetic changes which could be correlated with shifts in geography and climate, thereby facilitating understanding of local adaptation in the species. For example, identification of new locus contributing to phenotypic differences was discovered recently in snapdragons by whole-genome cline scans in hybrid zones (Field et al., 2026).

FR-enriched light is perceived as shade by the trees, e.g. under canopy, most of the R light is absorbed by the plant pigments while FR is reflected leading to reduction in the R:FR ratio (Ballare et al., 1987). A low R:FR ratio triggers the plant’s shade avoidance response – the survival mechanism by altering the growth patterns (Ranade et al., 2022, Ranade et al., 2019). Light quality varies with latitude across Sweden; the ratio of red (R) to far-red (FR) light (R:FR) decreases as you move from south to north, leading to an increased amount of FR (twilight) in the northern latitudes which is similar to shade-like condition (Nilsen, 1985). This gradient shapes how native flora including local Norway spruce populations regulate seasonal growth and flowering during the extended daylight periods comprising longer exposure to FR light in the northern latitudes during the growing season. Norway spruce exhibits a cline for growth in response to light quality in the Swedish populations (Clapham et al., 1998, Molmann et al., 2006, Ranade and García-Gil, 2021).

Norway spruce is a shade tolerant conifer species, meaning it can establish and compete well under shaded conditions (Ranade and García-Gil, 2023, Grebner et al., 2021, Ranade et al., 2022, Ranade et al., 2019). The species shows a requirement for FR light to maintain growth that follows a latitudinal gradient (Clapham et al., 1998, Molmann et al., 2006, Ranade and García-Gil, 2021). Norway spruce has evolved distinct genetic variations that follow a latitudinal cline in several genes e.g. *phytochrome*, *cryptochrome*, *expansin B3* and *MYB domain protein 3* that reflects local adaptation in the species (Ranade and García-Gil, 2023, Ranade and García-Gil, 2021). For example, cSNP variations in phytochromes which are the central regulators of the light signalling pathway, interpret regional differences in far-red signals, thus adapting to the local light conditions across Sweden. Lignin synthesis and growth are influenced by light quality; Norway spruce displays latitudinal variation in cSNPs in the genes from lignin biosynthesis pathway and variation in their expression along with differential lignin formation, influencing disease resilience (Ranade et al., 2022). Northern populations of Norway spruce are slow growing but disease resilient as compared to the southern populations; the northern trees have enhanced lignin synthesis and expression of higher number of defence-related genes during growth seasons in response to the prevailing light conditions namely higher exposure periods to FR light (Ranade et al., 2022). Phenological traits like bud-burst and bud-set driven by environmental cues like temperature also show a strong latitudinal cline for bud phenology in Norway spruce (Chen et al., 2014, Skroppa et al., 2010, Mikola, 1982). The northern populations of Norway spruce have adapted to shorter growing seasons by initiating bud-set earlier and requiring less accumulated heat (lower temperature) to trigger spring bud-burst than southern populations.

Understanding clines, particularly involving local adaptation is challenging yet significant for managing long-lived species like Norway spruce that has widespread distributions and is economically important species for the forest industry in Sweden. Related information in this domain could be used in tree breeding methods to design assisted migration strategies as trees are genetically programmed for specific growing season length, frost tolerance and light quality. This can facilitate selection of locally adapted genotypes that optimise the balance between high survival rates matching the changing climate and sustained productivity, for breeding programs. In addition, it can also facilitate selection of families resistant to region-specific diseases or pathogen resilient varieties in general.

The current investigation aims at genome-wide identification of clinal variation in genes in Norway spruce populations across Sweden with the exome capture dataset, to explore the genetic and molecular basis of clinal adaptations. The study includes exome capture data from 1654 unrelated Norway spruce trees sampled across a wide latitudinal range in Sweden (55-67°N) that has variation in the photoperiod and temperature during the growth season. This data is most ideal to examine the SNP variation in a series of genes that may be correlated to the variation in the environmental/climatic parameters.

## Material and Methods

### Plant material

1654 individuals from natural forests (with unrelated parents), from different latitudes across Sweden were recruited in this study (Ranade and García-Gil, 2023). Individuals were divided into six populations considering the same principle described in Ranade and García-Gil, 2023 that did not follow any specific strategy, however it ensured that all the groups were continuous populations allowing free pollen exchange (Ranade and García-Gil, 2023). Six populations were represented by groups S1-S6. S1 comprised 245 trees from latitudes 55-57°N, S2 – 213 trees from latitude 58°N, S3 – 187 trees from latitudes 59-60°N, S4 – 213 trees from latitudes 61-62°N, S5 – 573 trees from latitudes 63-64°N and S6 – 223 trees from latitudes 65-67°N. These individuals represent a subset of a study that reports the structure of Swedish populations of Norway Spruce, showing two distinct groups – one includes central and southern populations in Sweden, while the other includes the populations from northern Sweden (Chen et al., 2021).

### SNP detection, diversity and detection of latitudinal cline

Norway spruce genome (v1.0) genome retrieved from the PlantGenIE resource (Sundell et al., 2015) was used as the reference genome for this analysis. SNP variation was derived by using exome capture, described in Baison et al. (Baison et al., 2019). Only bi-allelic missense cSNPs i.e. missense SNP from coding regions, were considered for analysis in this study. Missense cSNP represents a change in the protein-coding gene resulting in the substitution of one amino acid to another. This alteration in the amino acid leads to a change in the sequence of the encoded protein, which can impact its primary structure, protein folding, and stability, potentially leading to modification in its mode of function. The change in the amino acid possibly has a greater effect if it involves a change from hydrophilic to hydrophobic amino acid or polar to non-polar and vice-versa. Filtering criteria applied to VCF files was as follows, -- min-alleles 2 --max-alleles 2 --maf 0.01 --remove-indels --minQ 10 --max-missing 0.9.

Allele and genotype frequencies across the six populations were determined using SNPassoc (Gonzalez et al., 2007) and analysis of variance and Tukey’s posthoc tests (Bonferroni *p values*) were applied to determine the statistical significance of their differences. When there was a latitude-wise consecutive increase/decrease in allele frequency across the six populations, the allele was documented to exhibit a cline. The steepness of cline in the given allele was calculated as the difference in allele frequency between the southern-most and the northern-most populations, e.g. 0.6 was the steepest cline detected in this study, while 0.1 was considered to be the milder one. Latitudinal cline was categorised as standard cline and trend cline. In a standard cline, the allele frequencies either increased or decreased in a steady manner with each population from south to north, e.g. in case of the cSNPs with steepest clines (Figure 1 and Figure 2). The trend cline either shows an increasing trend or decreasing trend in the allele frequencies in the populations from south to north, meaning the increase or decrease in the allele frequencies does not occur serially or latitude wise, instead it shows a trendline. For example, a cSNP (Thr125Ala, Polar hydrophilic > Non-polar hydrophobic) from LHCB7 shows high cline steepness of 0.4; however, although populations 1 to 5 show a steady increase in the allele frequency, there is a decrease in allele frequency in population 6 as compared to the population 5 (Supplementary file1 – Table S1; Supplementary file2 – Figure S21). Genetic diversity among the six Norway spruce populations across Sweden, was determined by Weir and Cockerham F_ST_ estimates. This included 238 512 biallelic missense SNPs from 1654 Norway spruce trees.

**Figure 1.**
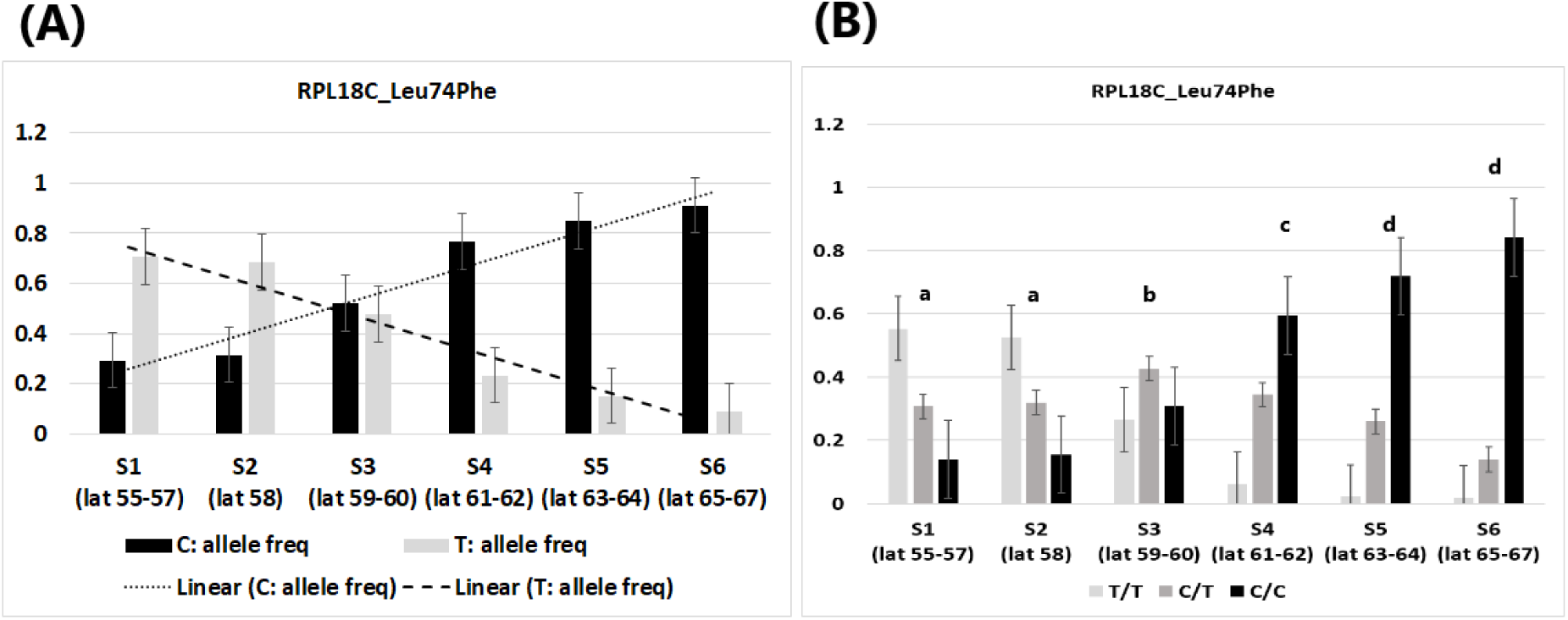
Allele and genotype frequencies of SNP variation in RPL18C gene showing cline across Norway spruce populations in Sweden. (a) Allele frequencies of Leu74Phe. (b) Genotype frequencies of Leu74Phe. One-way ANOVA and Tukey’s post-hoc test with the genotype frequencies showed significant differences across populations; Tukey’s post-hoc categorization is indicated above the bars.

**Figure 2.**
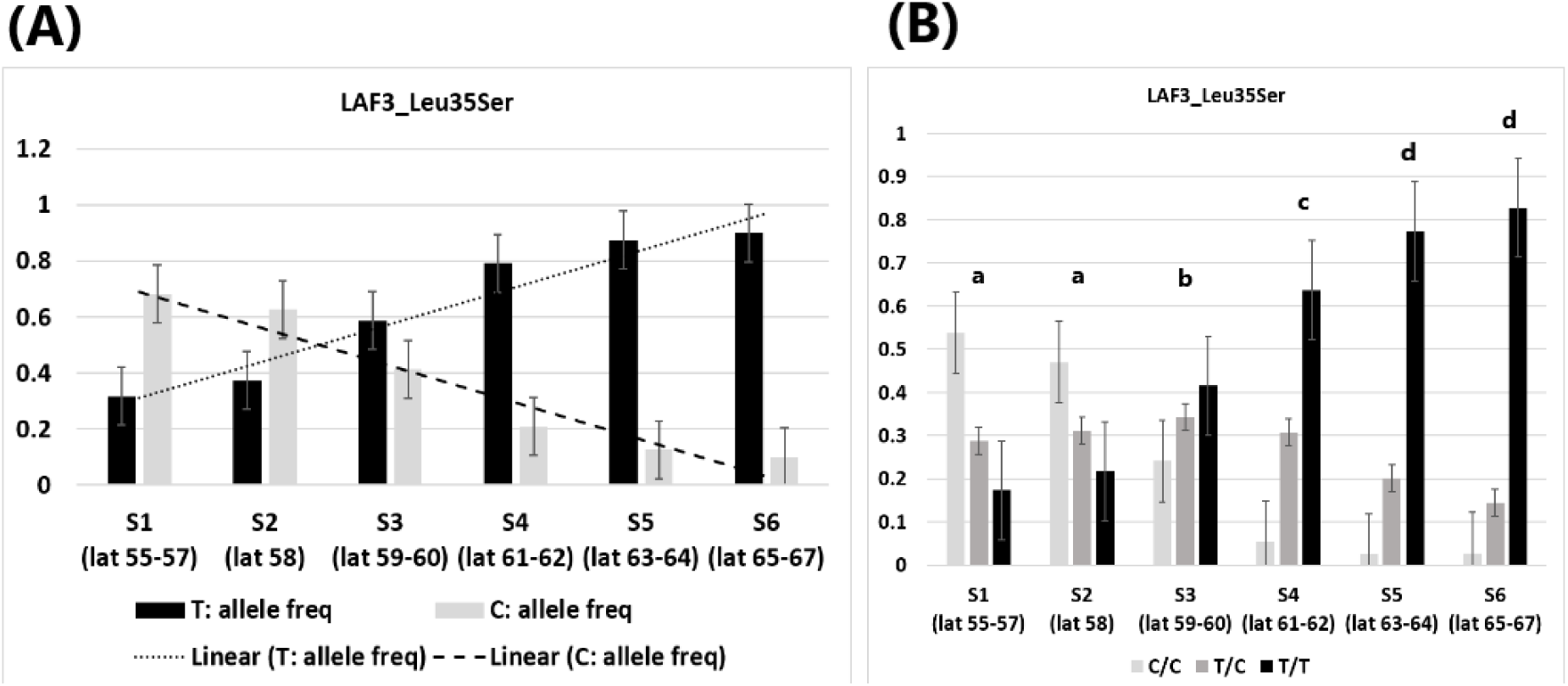
Allele and genotype frequencies of SNP variation in LAF3 gene showing cline across Norway spruce populations in Sweden. (a) Allele frequencies of Leu35Ser. (b) Genotype frequencies of Leu35Ser. One-way ANOVA and Tukey’s post-hoc test with the genotype frequencies showed significant differences across populations; Tukey’s post-hoc categorization is indicated above the bars.

### Gene ontology (GO) enrichment analysis

Gene ontology (GO) enrichment analysis was performed by using ShinyGO 0.85.1, with a p-value cut-off of 0.05 (FDR), using the *Arabidopsis* database, as *Arabidopsis* is the most well studied plant model (Ranade et al., 2022). GO:0003700 (DNA-binding transcription factor activity) was referred for deriving the transcription factors and they were grouped into types using PlantTFDB (Jin et al., 2017).

## Results

The filtering of the raw VCF file retained 238 512 biallelic SNPs. 37 912 missense cSNPs were annotated (PlantGenIE resource (Sundell et al., 2015)) having a representative *Arabidopsis* gene, representing 10161 MA ids (generally, one MA id represents one gene). Finally, 2935 missense cSNPs from the 1654 individuals across six populations, showed standard latitudinal cline in their allele/genotype frequencies. These SNPs belonged to 2084 annotated genes from Norway spruce including flowering and defence related genes. Genes including missense cSNPs with the steepest standard clines are included in Table 1 and Table 2, while a complete list of all the missense cSNPs showing standard and trend cline is represented in the Supplementary file1 – Table S1. Alignments of the Norway spruce genes with the model plant *Arabidopsis* for the genes that show steepest clines are included in Supplementary file2 (Figure S1-S20). The alignments file includes the details of the variation that shows standard cline e.g. nucleotide/codon/amino acid alteration and the cline steepness.

**Table 1.** Genes showing steep latitudinal cline in cSNPs in Norway spruce populations across Sweden primarily belong to defence/DNA repair pathways. Change in the amino acid property e.g. polar to non-polar/ hydrophilic to hydrophobic and vice-versa is indicated in bold.

| Gene name | Spruce gene ID (PlantGenIE)/Arabidopsis gene ID (TAIR) | Variation/Cline steepness | Function | Reference |
|---|---|---|---|---|
| *60S RIBOSOMAL PROTEIN L18* (*RPL18C*) | MA_3674g0020/ AT5G27850 | CTC > TTC, Leu74Phe<br>Non-polar hydrophobic > Non-polar hydrophobic<br>Cline steepness: 0.6 | Component of 60S subunit, facilitates ribosome assembly and translation, plant survival and defence/stress response. | (Tao et al., 2024) |
| *LONG AFTER FAR-RED 3* (*LAF3*) | MA_9446113g0010/ AT3G55850 | TTA > TCA, Leu35Ser<br>**Non-polar hydrophobic > Polar hydrophilic**<br>Cline steepness: 0.6 | Mediates Phytochrome A dependent responses under far-red light e.g. hypocotyl growth inhibition. | (Hare et al., 2003) |
| *ZINC 4 FINGER DNA 3′-PHOSPHOESTERASE* (*ZDP*) | MA_39122g0010/ AT3G14890 | CCA > CGA, Pro22Arg<br>**Non-polar hydrophobic > Polar hydrophilic**<br>Cline steepness: 0.5 | Involved in DNA repair and DNA demethylation pathways, maintenance of epigenome and genome stability. | (Li et al., 2025) |
|  |  | AAC > GAC, Asn31Asp<br>Polar hydrophilic > Polar hydrophilic<br>Cline steepness: 0.5 |  |  |
|  |  | TGT > TAT, Cys69Tyr<br>Polar hydrophilic > Polar hydrophilic<br>Cline steepness: 0.5 |  |  |
| *COENZYME Q9* (*COQ9*) | MA_9293920g0010/ AT1G19140 | TCT > GCT, Ser77Ala<br>**Polar hydrophilic > Non-polar hydrophobic**<br>Cline steepness: 0.5 | Involved electron transfer respiratory chain crucial for ATP production, in mitochondria. Acts as antioxidant. | (Xu et al., 2025) |
| *SERINE/ARGININE RICH-LIKE PROTEIN 45A* (*SR45A*) | MA_42236g0030/ AT1G07350 | AGC > ATC, Ser209Ile<br>**Polar hydrophilic > Non-polar hydrophobic**<br>Cline steepness: 0.5 | Abiotic stress - regulator of tolerance to heat, salinity and drought. | (Muthusamy et al., 2020) |
| *ATP-BINDING CASSETTE G26* (*ABCG26*) | MA_16472g0020/ AT3G13220 | GAA > GAT, Glu34Asp<br>Polar hydrophilic > Polar hydrophilic<br>Cline steepness: 0.5 | Essential for pollen development and male fertility. | (Zhao et al., 2016) |
| *OXIDATION RESISTANCE 1* (*OXR1*) | MA_697287g0010/ AT4G39870 | TCA > GCA, Ser129Ala<br>**Polar hydrophilic > Non-polar hydrophobic**<br>Cline steepness: 0.5 | Offers protection against oxidative stress and maintains genome stability. | (Colombatti et al., 2019) |
| *ALUMINUM-ACTIVATED MALATE TRANSPORTER 9* (*ALMT9*) | MA_10243484g0010/ AT3G18440 | GTA > TTA, Val122Leu<br>Non-polar hydrophobic > Non-polar hydrophobic<br>Cline steepness: 0.5 | Regulation of stomatal opening and osmotic balance. | (De Angeli et al., 2013) |
| *TERPENE SYNTHASE 02* (*TPS02*) | MA_10426551g0010/ AT4G16730 | TGG > TTG, Trp43Leu<br>Non-polar hydrophobic > Non-polar hydrophobic<br>Cline steepness: 0.5 | Defence against pathogens, insects, herbivores. | (Huang et al., 2010) |

**Table 2.** Genes primarily involved in flowering showing steepest latitudinal cline in cSNPs in Norway spruce populations across Sweden. Change in the amino acid property e.g. polar to non-polar/ hydrophilic to hydrophobic and vice-versa is indicated in bold.

| **Gene name**<br>**\*TF (Transcription Factor)** | **Spruce gene ID (PlantGenIE)/Arabidopsis gene ID (TAIR)** | **Variation/Cline steepness** | **Function** | **Reference** |
|---|---|---|---|---|
| *BARELY ANY MERISTEM 1* (*BAM1*) | MA_40935g0020/AT5G65700 | TAT > TCT, Tyr158Ser<br>Polar hydrophilic > Polar hydrophilic<br>Cline steepness: 0.4 | Required for floral meristem development, promotes anther development. | (Hord et al., 2006, DeYoung et al., 2006) |
|  |  | GGC > GAC, Gly174Asp<br>**Non-polar hydrophobic > Polar hydrophilic**<br>Cline steepness: 0.4 |  |  |
| *REDUCED VERNALIZATION RESPONSE 1* (*VRN1*)<br>**\*TF type: B3** | MA_10430774g0010/AT3G18990 | TCA> TTA, Ser641Leu<br>**Polar hydrophilic > Non-polar hydrophobic**<br>Cline steepness: 0.4 | Floral activator, involved in transition from vegetative growth to reproductive development. | (King et al., 2013) |
| *JMJC DOMAIN-CONTAINING PROTEIN 27* (*JMJ27*) | MA_115863g0010/AT4G00990 | ACT > GCT, Thr1103Ala<br>**Polar hydrophilic > Non-polar hydrophobic**<br>Cline steepness: 0.3 | Repressor of flowering and positive modulator of immune responses | (Dutta et al., 2017) |
| *LATE MERISTEM IDENTITY2* (*LMI2*)<br>**\* TF type: R2R3 MYB** | MA_128244g0010/AT3G61250 | TGT > AGT, Cys217Ser<br>Polar hydrophilic > Polar hydrophilic<br>Cline steepness: 0.3 | Promotes transition from vegetative growth to flowering. | (Pastore et al., 2011) |
|  | MA_118448g0010/AT3G61250 | GGG > AGG, Gly140Arg<br>**Non-polar hydrophobic > Polar hydrophilic**<br>Cline steepness: 0.3 |  |  |
| *FLOWERING CONTROL LOCUS A* (*FCA*) | MA_5232462g0010/AT4G16280 | CAA > AAA, Gln392Lys<br>Polar hydrophilic > Polar hydrophilic<br>Cline steepness: 0.2 | Promotes transition from vegetative growth to flowering. | (Macknight et al., 1997) |
| *TERMINAL FLOWER 1* (*TFL1*) | MA_178498g0010/AT5G03840 | GAT > GAA, Asp54Glu<br>Polar hydrophilic > Polar hydrophilic<br>Cline steepness: 0.2 | Suppresses floral meristem, involved in the regulation of flowering time. | (Hanano and Goto, 2011) |

Cline steepness in the allele frequencies of an allele was calculated as the difference in allele frequency between the southern-most and the northern-most populations. Steepest standard clines in the allele frequencies of missense cSNPs were detected in *60S RIBOSOMAL PROTEIN L18* (*RPL18C*), a gene involved in plant survival and defence/stress responses (Figure 1) and *LONG AFTER FAR-RED 3* (*LAF3*) that mediates light responses (Figure 2). This was closely followed by cline in *ZINC 4 FINGER DNA 3’-PHOSPHOESTERASE* (*ZDP*), a gene involved in DNA repair and DNA demethylation pathways (Table 1), which showed cline in three cSNPs. cSNPs from genes involved in ATP production (*COENZYME Q9*; *COQ9*), regulation of abiotic stress (*SERINE/ARGININE RICH-LIKE PROTEIN 45A*; *SR45A*), oxidative stress (*OXIDATION RESISTANCE 1*; *OXR1*), osmotic balance (*ALUMINUM-ACTIVATED MALATE TRANSPORTER 9*; *ALMT9*), pollen development (*ATP-BINDING CASSETTE G26*; *ABCG26*) and defence response (*TERPENE SYNTHASE 02*; *TPS02*) also exhibited steep clines (Table 1). The allele frequencies, genotype frequencies and the p-values for Hardy-Weinberg Equilibrium (HWE) from the missense cSNPs with steepest clines are included in Supplementary file1 – Table S2.

Genes from reproductive pathways involved in flowering that showed the steepest clines in allele frequencies of missense cSNPs included *BARELY ANY MERISTEM 1* (*BAM1*), *REDUCED VERNALIZATION RESPONSE 1* (*VRN1*), closely followed by *JMJC DOMAIN-CONTAINING PROTEIN 27* (*JMJ27*) and *LATE MERISTEM IDENTITY2* (*LMI2*) (Table 2). *FLOWERING CONTROL LOCUS A* (*FCA*) and *TERMINAL FLOWER 1* (*TFL1*) involved in transition from vegetative growth to flowering and regulation of flowering time, also displayed significant clines (Table 2). The allele frequencies, genotype frequencies and the p-values for HWE from the missense cSNPs from flowering genes are included in Supplementary file1 – Table S3. Approximately 50% of cSNPs with steep clines (Table 1, Table 2) bear a change in the amino acid from hydrophilic to hydrophobic amino acid or polar to non-polar and vice-versa, indicating modification in the mode of function of the encoded protein.

The available version of Norway spruce genome (v1.0) during the analysis (Sundell et al., 2015), was fragmented, yet the sequence alignments (Supplementary file2 – Figure S1-S20) and tools applied on the PlantGenIE resource (Sundell et al., 2015) to identity the Norway spruce gene, represent consistent results. Due to fragmentation, the gene sequences are not complete, and some genes are proposed to be contained across two separate contigs. For example, the genes *RPL18C*, *LAF3* and *COQ9* are contained across two separate contigs (Supplementary file2 – Figure S1-S4, Figure S6-S7). Though, these could also potentially be the homologues of RPL18C, LAF3 and *COQ9* owing to fragmented genome. In any case, these gene fragments contain variations that show cline. Few of the genes show cline in multiple cSNPs; some cSNPs display steep cline while others represent milder clines. *LAF3* shows two missense variants with a cline steepness of 0.6 and 0.4, while and *COQ9* shows one missense variant with cline steepness of 0.5 and four other variants with cline steepness of 0.1 (Table1, Supplementary file2 – Figure S3-S4, Figure S6-S7). *SR45a*, *OXR1* and *TPS02* contain missense variants with cline steepness of 0.1, 0.4 and 0.3, respectively (Table1, Supplementary file1 – Table S1; Supplementary file2 – Figure S8, Figure S10, Figure S12). Likewise, with reference to flowering genes (Table 2) *FCA* has four additional missense variants, each with a cline steepness of 0.1 while homologues of *LMI2* (Supplementary file2 – Figure S16-S18), *AGAMOUS-LIKE 20* (*AGL20*) and *Apetala 2* (*AP2*) have variants representing mild (0.1) to very mild cline (<0.1) (Supplementary file1 – Table S1).

Gene ontology (GO) enrichment analysis was conducted with 2084 genes that were annotated in Norway spruce, harbouring 2935 missense cSNPs from 1654 individuals across the six populations included in this study. GO analysis for Biological Process indicated higher fold enrichment for developmental processes involved in reproduction, along with response to hormone, endogenous stimulus and processes related to post-embryonic development, which can be correlated to the clines detected in the flowering genes (Supplementary file2 – Figure S22). GO analysis for Molecular Function showed significantly higher representation of genes involved in nucleic acid binding indicating active transcription and DNA repair mechanisms (Supplementary file2 – Figure S23). This coincides with one of the steepest clines detected in *ZDP*, a gene involved in DNA repair. Cellular Compartment GO analysis included higher representation of genes localised in the plastids, while higher fold enrichment was represented by nuclear body category (Supplementary file2 – Figure S24).

MYB family members represented the highest numbers among all the transcription factors that followed a cline in the Norway spruce populations included in this study (Figure 3). MYBs are primarily involved in secondary cell wall formation, lignin deposition, wood development and stress response (Wu et al., 2024, Ranade and García-Gil, 2024). MYBs were closely followed by EFRs and bHLH family members. ERFs act downstream of the ethylene signalling pathway in addition to response to stress and defence (Wu et al., 2022) while bHLHs are involved in light signalling, regulation of growth and development (Zuo et al., 2023).

**Figure 3.**
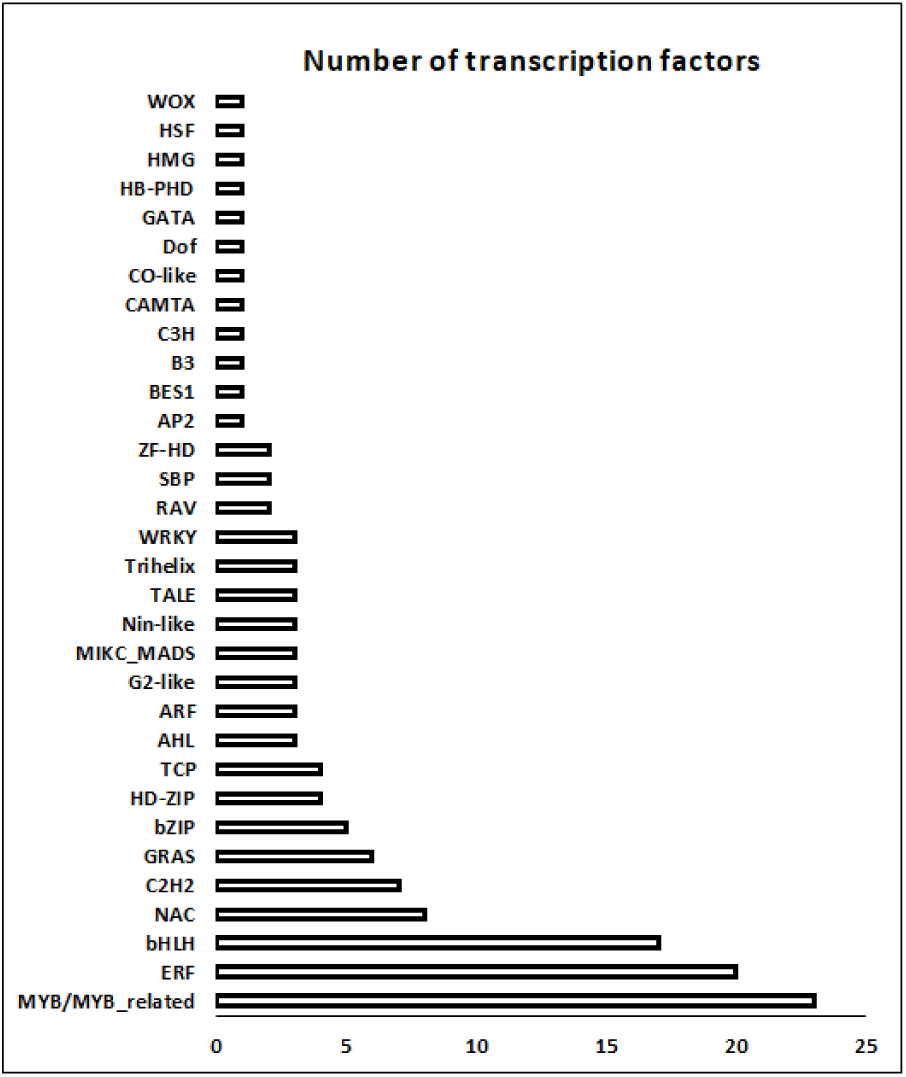
Number of transcription factors of each type, with missense cSNPs showing clinal variation across Norway spruce populations in Sweden.

Weir and Cockerham FST estimates for six populations of Norway spruce across Sweden showed low values (Table 3).

**Table 3.**
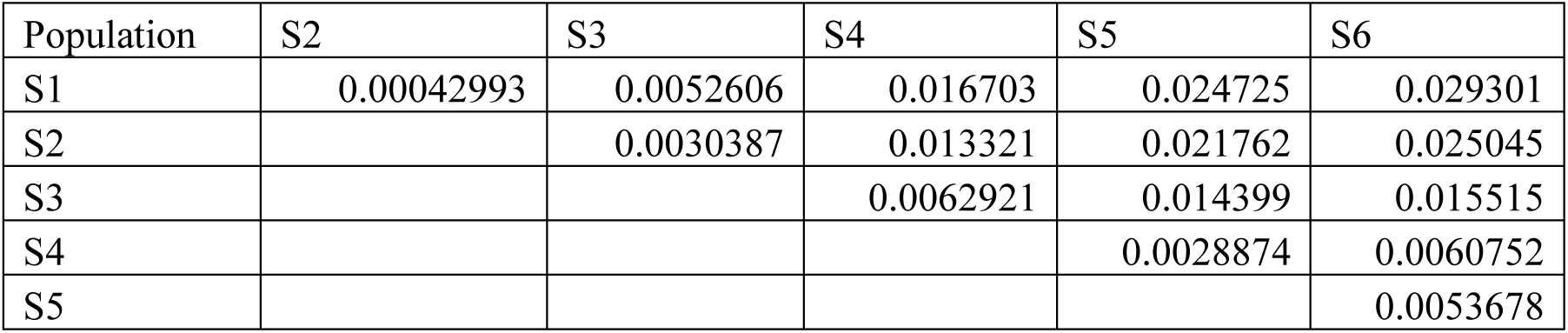
Weir and Cockerham F_ST_ estimates for six populations (1654 Individuals) of Norway spruce across Sweden involved in the detection of latitudinal clines (including 238 512 biallelic missense SNPs).

## Discussion

### Weak neutral genetic differentiation contrasts with steep adaptive trait clines following postglacial recolonisation

Conifers originated approximately 300 million years ago and they have evolved over millions of years to adapt to steep latitudinal and longitudinal environmental gradients (Williams, 2009). The genetic composition of Norway spruce populations in Sweden reflects this evolutionary history, which underwent post-glacial recolonisation by spruce migrating from two distinct Last Glacial Maximum (LGM) refugia. Migration from the east via Russia and Finland contributed to the recolonisation of Scandinavia, while Central European ancestry also contributed to the genetic composition of southern and central Swedish populations (Chen et al., 2019).

Norway spruce populations in Sweden appear to show weak population structure, with low neutral genetic differentiation between populations (F_ST_) owing to large population sizes and a continuous distribution with few geographic barriers, allowing high pollen-mediated gene flow (Chen et al., 2021, Eklöf et al., 2026). The Swedish Norway spruce population comprises two distinct groups; the central and southern populations form one group and the other includes the populations from northern Sweden (Chen et al., 2021). Weir and Cockerham F_ST_ estimates for six populations of Norway spruce across Sweden (Table 3) agree with several previous findings regarding low F_ST_ values suggesting low genetic differentiation (Chen et al., 2016, Ranade and García-Gil, 2021, Ranade and García-Gil, 2023, Wang et al., 2020, Androsiuk et al., 2013).

A weak neutral population differentiation contrasts with a strong cline for adaptive traits (Q_ST_). The strongest evidence for local adaptation and optimal survival under given environmental conditions in Norway spruce populations in Sweden is provided by the traits such as timing of bud-burst and bud-set, as well as cold hardiness, height and drought tolerance (Clapham et al., 1998, Sogaard et al., 2008, Westin et al., 2000a, Westin et al., 2000b, Notivol et al., 2007). Adaptive clines are the result of adaptation to temperature and daylength; in addition, light quality also plays a crucial role (Ranade and García-Gil, 2021, Ranade and García-Gil, 2023, Westin et al., 2000a). The northern populations of Norway spruce require far-red enriched light to maintain growth, cease growth earlier, are more frost-resistant and disease-resilient than southern populations, indicating adaptation to local climatic conditions (Ranade et al., 2022, Ranade and García-Gil, 2021, Molmann et al., 2006, Westin et al., 2000b).

The genetic basis of the steep clines observed for adaptive traits are the result of clinal variation in the DNA sequence and expression patterns of genes involved in phenology, lignin synthesis and defence ultimately influencing trait variation along the cline (Ranade and García-Gil, 2021, Ranade and García-Gil, 2023, Ranade and García-Gil, 2025, Ranade et al., 2022). The current work provides further support for local adaptation in Norway spruce in genes regulating DNA repair, defence response and flowering. Previously, the north-south genetic gradients in allele frequency and gene expression have been detected in several genes including phytochromes (central regulators of light pathway), genes involved in cell wall formation and defence, which are proposed to be due to local environmental adaptation (Ranade and García-Gil, 2021, Ranade and García-Gil, 2023, Ranade et al., 2022). In those cases, F_ST_ values increase with geographic distance, potentially reflecting reduced pollen-mediated gene flow after certain distance and local adaptation to environmental conditions that favour different alleles.

### Latitudinal clines in the regulation of flowering

Reproduction is fundamental for species survival. Adaptation and genetic diversity plays a crucial role in the process of reproduction. Flowering and seed production are the key events in the reproductive cycle in plants. The flowering phase involves a shift from vegetative to reproductive growth, initiating a reallocation of resources, often affecting the tree biomass, with effects depending on the species and environmental conditions (Zhao et al., 2021). Flowering depends on the climatic conditions; plants flower early or late depending mainly on the light conditions (photoperiod), temperature and degree days. Recent studies in flowering plants have reported that initiation of flowering correlates with latitude as the process is governed by climatic gradients (Landoni et al., 2024). Ecotypic variation in flowering time owing to sequence variations in flowering genes is associated with adaptation across natural seasonal environments (Fournier-Level et al., 2022). Furthermore, epistasis can influence flowering time following a latitudinal cline (Caicedo et al., 2004). However, in conifers this area remains less explored from the molecular point of view. The current work may provide the basis for understanding this complex network as we report gene variation in flowering-related genes that may contribute to difference in flowering time in different Norway spruce populations; e.g. the northern Norway spruce populations flower early in the growing season than the southern while the southern ones flower more frequently compared to the northern populations (Danusevicius and Brazaitis, 2011, Nikkanen and Ruotsalainen, 2000). In the current analysis, moderate to mild clinal variations in cSNPs were detected in floral activators (*VRN1*, *LMI2*, *FCA*) as well as repressors of flowering (*JMJ27*, *TFL1*), along with genes that regulate the development of the floral meristem in general (*BAM1*) (Table 2). *VRN1,* which showed a moderate cline of steepness of 0.4, encodes a B3-type of TF essential for the repression of FLC in vernalized in *Arabidopsis* plants (King et al., 2013). Other target genes of *VRN1* includes genes that regulate frost tolerance (Deng et al., 2015). This is consistent with the latitudinal temperature gradient in Sweden, where northern latitudes experience much lower freezing temperatures than southern latitudes. SNP variation along the temperature gradient might have resulted in populations adapted to the local temperature conditions.

Although the scope of the current work was limited only to missense mutations, it is worth analysing the synonymous polymorphisms because they can alter RNA secondary structure. Clinal variation was detected in a synonymous cSNP (771C>T) in *SQUAMOSA BINDING PROTEIN-LIKE1* (*SPL1*) gene that harbours the *acrocona* mutant-specific SNP in its miRNA156/ miRNA529 binding site due to which this mutant displays early cone setting (Akhter et al., 2022). *SPL1* is active in reproductive meristems while miRNA156/miRNA529 are present in high levels at juvenile plant stages and they prevent early flowering by suppressing the *SPL1* gene. The *acrocona* mutant-specific SNP reduces miRNA156/miRNA529-mediated repression, allowing early cone setting (Akhter et al., 2022). A synonymous polymorphism can modulate microRNA(miRNA)-mRNA interactions and can and alter the secondary RNA structure which may influence the accessibility of miRNA-binding sites. The 771C>T SNP from *SPL1* displays a standard cline and it is present 309 bp away from the miR529/miR156-binding site (Supplementary file2 – Figure S25). Although this SNP is located too far (309 bp) to directly disrupt the binding site, it can still affect gene regulation by changing the overall structure of the RNA strand and influence the availability of miRNA-binding sites (Kertesz et al., 2007). In this context, the 771C>T SNP from *SPL1*, may be involved in the regulation of flowering time. With the given scenario, it is noteworthy that the data from the current investigation come from exome capture and hence do not cover the entire gene. It would be strategic to sequence entire *SPL1* gene from a sufficiently large population along the latitudinal cline, to reveal other SNPs located in the vicinity of miRNA156/ miRNA529-binding site, which would give information of one of the key aspects regarding molecular basis of regulation of flowering time in Norway spruce.

The current analysis was performed using an exome capture dataset; therefore, the SNPs were detected from a small sequence from the coding region of the gene. Although a synonymous SNP does not change the encoded amino acid, yet it identifies a variable region. A missense SNP positioned close to a synonymous SNP may have remained unidentified because the exome capture probe was not designed to detect it. The neighbouring SNPs are often in linkage disequilibrium and may show similar allele frequencies and association signals. In this context, yet another synonymous SNP from *STYLISH1* (*STY1*) displayed a steep cline of 0.6 (Supplementary file2 – Figure S26); *STY1* is a transcription factor that regulates floral organ development and plays a role in controlling flowering time as well as in the maintenance of apical meristem (Ståldal et al., 2012). Sequencing the entire gene will provide a more comprehensive picture of local adaptation associated with the gene of interest.

Variants from several transcription factors involved in the regulation of flowering displayed clinal variation. Few examples of key TFs include *VRN1* (King et al., 2013), *LMI2* (Pastore et al., 2011), *AGAMOUS-LIKE 20* (*AGL20*), *APETALA 2 (AP2*) (Yant et al., 2010) and *LONG VEGETATIVE PHASE 1* (*LOV1*) (Yoo et al., 2007). These TFs have cline steepness values ranging from 0.1 to 0.4 and belong to the B3, R2R3 MYB, MADS box, AP2/EREBP and NAC-domain transcription factor types, respectively (Supplementary file1 – Table S1). All these TFs are key regulators of flowering and control the formation of floral organs and flowering time. In addition to repression of flowering, *LOV1* plays a central role in promoting cold/freezing tolerance (Yoo et al., 2007) and it was found to be upregulated in response to FR-enriched light in the northern Norway spruce populations as compared to the southern ones (Ranade and García-Gil, 2021). Here again it is worth noticing the steep temperature gradient along the latitudes under consideration. AGL20 acts as a major floral activator and it promotes the transition to flowering (Lee et al., 2000), while AP2 represses flowering; AP2 acts as a repressor to inhibit AGL20 (Yant et al., 2010). Overall, regulation of flowering involves a complex network of TFs and other genes involved in the formation of floral organs which are interconnected. cSNPs variants that follow a latitudinal cline represent local adaptations to the climatic clues and facilitate the tree to survive, thrive and reproduce at its best in its environment.

### Association of variation in cSNPs with bud-set/bud-burst cline

Bud-set or growth cessation occurs in late summer/autumn, and bud-burst refers to growth reactivation in spring. Both traits follow steep latitudinal clines in Norway spruce and depend on climatic cues, namely photoperiod and temperature; they allow the tree to maximize its growing season while minimising the risk of severe frost damage (Svystun et al., 2021, Ranade and García-Gil, 2021). Bud-burst is governed mainly by temperature; the northern Norway spruce populations flush earlier with the first signs of sustained warmth, while southern trees result in later bud-burst that protects them against spring frost (Sogaard et al., 2008, Svystun et al., 2021). Epigenetic memory plays a crucial role in Norway spruce growth rhythm, for example, epigenetic memory of warm temperatures during embryogenesis leads to earlier bud-burst altering the expression of key phenological genes like *FLOWERING LOCUS T/TERMINAL FLOWER 1-*like gene (*PaFTL2*) (Carneros et al., 2017). In Norway spruce, the *Terminal Flower 1* and *Terminal Flower 2* refer to FLOWERING LOCUS T/TERMINAL FLOWER 1-Like genes (abbreviated as *PaFTL1* and *PaFTL2*) (Karlgren et al., 2013). *PaFTL1* and *PaFTL2* are paralogous genes; their temporal and spatial expression largely complement each other, suggesting their simultaneous role in regulating perennial growth in Norway spruce (Karlgren et al., 2013). While *TFL1* represses floral meristem development, *TFL2* induces bud-set and growth cessation in *Arabidopsis* (Larsson et al., 1998). Although the timing of bud-set and bud-burst following a steep latitudinal cline has been well documented in Norway spruce, the underlying molecular mechanisms are less explored owing to the complex multigenic involvement. In the current analysis, we detected a moderate cline in a cSNP associated with *TFL1* or the *PaFTL1*, which may contribute to clinal variation of bud-set/bud-burst in Norway spruce.

### Stress response and disease resistance

*RPL18C, ZDP, OXR1* and *TPS02* (Table 1) are defence-related genes whose cSNPs show steep clines in allele frequency. *RPL18C* plays a key role in the defence mechanism in eukaryotes including humans (Dhar et al., 2006). Likewise, *OXR1* protects the cells from oxidative stress and mediates DNA repair in mammalian cells too (Matsui et al., 2020). Moreover, the tolerance to oxidative stress may lead to an increase in plant biomass (Colombatti et al., 2019). Interesting ecotypic variation has been observed in *Arabidopsis*; the Col-0 ecotype in *Arabidopsis* possesses a non-functional *TPS02* due to a frame-shift mutation, whereas in the Ws ecotype, it is a key functional gene without the mutation (Huang et al., 2010). In this context, the variation in Norway spruce *TPS02* may be linked to a similar phenomenon conferring resistance to the northern populations (Ranade et al., 2022). Three cSNPs were detected in *ZDP*, which is involved in DNA repair and defence; although the ZDP sequence obtained from PlantGenIE was partial, it contained the conserved protein domain HAD_PNP (Supplementary file2 – Figure S5) which was identified using the Conserved Domain Database (CDD); the HAD_PNP domain plays a key role in both base excision repair and non-homologous end-joining DNA repair pathways (Marchler-Bauer et al., 2015).

cSNPs from several genes (Table 1, Supplementary file1 – Table S1) involved in stress response and disease resistance followed a latitudinal cline including genes from various pathways contributing to disease resistance like the lignin, phenylpropanoid, terpenoid and flavonoid pathways (e.g. MYBs and peroxidases). Another example is MAP KINASE SUBSTRATE 1 (MKS1) which plays a key role in modulating plant innate immunity and regulating pathogen defence mechanisms through salicylic acid and jasmonate pathways (Andreasson et al., 2005). Several genes from the Leucine-Rich Repeat (LRR) family show clines in the cSNPs (Supplementary file1 – Table S1); LRR family proteins are immune receptors involved in plant defence (Hao et al., 2025). cSNPs from genes regulating drought stress responses, such as aquaporin genes (Afzal et al., 2016) and genes involved in abscisic acid (ABA) signalling (Sharma et al., 2023), display moderate clines. Although majority of these cSNPs displayed mild clines, the respective genes involved may have a combined effect on the defence mechanism, through complex networks taking part in the stress response.

### Genes contributing to “tree-ness” and tree growth in general

A recent comparative analysis of tree species proposed that unique genes defining a “woody tree”, do not exist. Instead, trees have a genetic makeup similar to that of non-woody plants and herbs. The study concluded that the trees organize and regulate these shared genes differently which renders them the “tree-ness”. Few examples of such genes are expansins, laccases, peroxidases and the MYB-related genes, which are essential for secondary cell wall development leading to wood formation (Supplementary file1 – Table S1) (Ranade et al., 2022, Ranade and García-Gil, 2025). Genes involved in plant growth in general, displayed moderate/mild clines in their cSNPs; few examples include *SLOW GROWTH* genes (*SLO2*, *SLO3*), *FCS LIKE ZINC FINGER 6* (*FLZ6*) and *FRUCTOSE-BISPHOSPHATE ALDOLASE 6* (*FBA6*) (Hsieh et al., 2015, Jamsheer et al., 2018, Liu et al., 2025) (Supplementary file1 – Table S1).

Light pathways modulate environmental signalling and are the key contributors towards metabolic energy production controlling the overall tree growth. It has already been documented that phytochromes, central regulators of light pathway show distinct clines in their allele frequencies (Ranade and García-Gil, 2023). Another worth mentioning example from this pathway is *LIGHT-HARVESTING COMPLEX B7* (*LHCB7*), a chlorophyll A-B binding family protein that fine-tunes the rate of photosynthesis (Mei et al., 2025). A missense cSNP (Thr125Ala) from *LHCB7* displayed a trend cline, having a cline steepness of 0.4 (Supplementary file2, Figure S21).

### Significance of clinal variation for tree breeding

Clinal variation provides a useful basis for selecting forest reproductive material suited to different growing conditions. In Norway spruce, traits such as cold hardiness, bud flush and growth cessation vary with latitude, reflecting adaptation to local temperature and light conditions. Linking these patterns to genetic variation can help identify material that combines growth with frost hardiness and disease resistance and guide its deployment as the climate changes. For example, northern Swedish populations generally show greater cold tolerance and have shown stronger defence responses, whereas southern populations tend to grow faster. Understanding the genetic basis of these differences can help breeders balance productivity and resilience, identify candidate markers for genomic selection, and guide assisted migration.

However, transferring trees along a latitudinal cline cannot be guided by temperature alone. Warming does not alter the daylength associated with a given latitude, and seasonal patterns of light quality will not simply shift northwards with temperature. Although light quality can vary with atmospheric and canopy conditions, its latitude-dependent seasonal patterns remain relevant to adaptation. Trees transferred northwards may therefore encounter warmer conditions that favour growth while responding inappropriately to local photoperiod and light quality, with consequences for growth cessation, bud set and cold acclimation. Such mismatches may also affect reproductive development, including the initiation and timing of flowering and the synchronisation of pollen release and female receptivity, potentially influencing seed production. Breeding and deployment strategies should therefore evaluate genetic material that combines favourable growth and defence alleles with alleles supporting vegetative and reproductive responses adapted to the light conditions at the site of deployment. Clinal information can help identify candidate combinations and define seed-transfer limits, reducing the risk of winter damage, poor growth or impaired reproductive performance when southern or coastal material is planted at northern or high-elevation sites (Luoranen et al., 2022).

## Conclusions

Our findings reveal pronounced clinal variation in genes associated with defence and reproduction, against a background of weak neutral genetic differentiation in Norway spruce. These genes provide candidates for understanding how populations adapt to contrasting growing conditions. The results also highlight why assisted migration, and breeding strategies should consider more than rising temperatures: daylength will remain unchanged, and seasonal light conditions will not shift northwards with warming. Evaluating genetic material that combines growth and defence with vegetative and reproductive responses suited to the destination could help reduce maladaptation and support productive, resilient forests under a changing climate.

## Acknowledgements

This work was partially supported by the Wallenberg Initiatives in Forest Research (WIFORCE) funded by the Knut and Alice Wallenberg Foundation (KAW 2020.0240). The authors acknowledge the support from the support from Science for Life Laboratory (SciLifeLab), the Swedish Foundation for Strategic Research (SSF), Bioinformatics Facility at the Umeå Plant Science Centre, the Uppsala Multidisciplinary Center for Advanced Computational Science at Uppsala University, and the PDC Center for High Performance Computing at KTH Royal Institute of Technology for access to computational resources. Computational resources were provided by the National Academic Infrastructure for Supercomputing in Sweden (NAISS), funded by the Swedish Research Council.

## Availability of data

The data that supports the findings of this study are available in the additional files of this article. The exome capture raw reads and the RNA-seq data have been deposited in NCBI’s sequence read archive (SRA) under accession number (PRJNA731384).

## Authors’ contributions

SSR: Conceptualization, Data curation, Formal analysis, Methodology, Software, Visualization, Writing – original draft, Writing – review & editing; MRGG: Conceptualization, Funding acquisition, Project administration, Resources, Supervision, Writing – review & editing.

## References

Afzal, Z., Howton, T. C., Sun, Y. L. & Mukhtar, M. S. 2016. The Roles of Aquaporins in Plant Stress Responses. Journal of Developmental Biology, 4.

Akhter, S., Westrin, K. J., Zivi, N., Nordal, V., Kretzschmar, W. W., Delhomme, N., Street, N. R., Nilsson, O., Emanuelsson, O. & Sundström, J. F. 2022. Cone-setting in spruce is regulated by conserved Elements of THE age-dependent flowering pathway. New Phytologist, 236, 1951–1963.

Andreasson, E., Jenkins, T., Brodersen, P., Thorgrimsen, S., Petersen, N. H. T., Zhu, S. J., Ǫiu, J. L., Micheelsen, P., Rocher, A., Petersen, M., Newman, M. A., Nielsen, H. B., Hirt, H., Somssich, I., Mattsson, O. & Mundy, J. 2005. The MAP kinase substrate MKS1 is a regulator of plant defense responses. Embo Journal, 24, 2579–2589.

Androsiuk, P., Shimono, A., Westin, J., Lindgren, D., Fries, A. & Wang, X. R. 2013. Genetic status of Norway spruce (*Picea abies*) breeding populations for northern Sweden. Silvae Genetica, 62, 127–136.

Baison, J., Vidalis, A., Zhou, L., Chen, Z.-Ǫ., Li, Z., Sillanpää, M. J., Bernhardsson, C., Scofield, D., Forsberg, N., Grahn, T., Olsson, L., Karlsson, B., Wu, H., Ingvarsson, P. K., Lundǫvist, S.-O., Niittylä, T. & García-Gil, M. R. 2019. Genome-wide association study identified novel candidate loci affecting wood formation in Norway spruce. The Plant Journal, 100, 83–100.

Ballare, C. L., Sanchez, R. A., Scopel, A. L., Casal, J. J. & Ghersa, C. M. 1987. Early detection of neighbor plants by phytochrome perception of spectral changes in reflected sunlight. Plant Cell and Environment, 10, 551–557.

Caicedo, A. L., Stinchcombe, J. R., Olsen, K. M., Schmitt, J. & Purugganan, M. D. 2004. Epistatic interaction between *Arabidopsis* FRI and FLC flowering time genes generates a latitudinal cline in a life history trait. Proceedings of the National Academy of Sciences of the United States of America, 101, 15670–15675.

Carneros, E., Yakovlev, I., Viejo, M., Olsen, J. E. & Fossdal, C. G. 2017. The epigenetic memory of temperature during embryogenesis modifies the expression of bud burst- related genes in Norway spruce epitypes. Planta, 246, 553–566.

Chen, J., Kallman, T., Ma, X. F., Zaina, G., Morgante, M. & Lascoux, M. 2016. Identifying genetic signatures of natural selection using pooled population sequencing in Picea abies. G3-Genes Genomes Genetics, 6, 1979–1989.

Chen, J., Li, L. L., Milesi, P., Jansson, G., Berlin, M., Karlsson, B., Aleksic, J., Vendramin, G. G. & Lascoux, M. 2019. Genomic data provide new insights on the demographic history and the extent of recent material transfers in Norway spruce. Evolutionary Applications, 12, 1539–1551.

Chen, J., Tsuda, Y., Stocks, M., Källman, T., Xu, N. N., Kärkkäinen, K., Huotari, T., Semerikov, V. L., Vendramin, G. G. & Lascoux, M. 2014. Clinal Variation at Phenology-Related Genes in Spruce: Parallel Evolution in FTL2 and Gigantea? Genetics, 197, 1025-+.

Chen, Z. Ǫ., Zan, Y. J., Milesi, P., Zhou, L. H., Chen, J., Li, L. L., Cui, B. B., Niu, S. H., Westin, J., Karlsson, B., GARCÍA-Gil, M. R., Lascoux, M. & Wu, H. X. 2021. Leveraging breeding programs and genomic data in Norway spruce (Picea abies L. Karst) for GWAS analysis (vol 22, 179, 2021). Genome Biology, 22.

Clapham, D. H., Dormling, I., Ekberg, I., Eriksson, G., Qamaruddin, M. & Vince-Prue, D. 1998. Latitudinal cline of requirement for far-red light for the photoperiodic control of budset and extension growth in *Picea abies* (Norway spruce). Physiologia Plantarum, 102, 71–78.

Colombatti, F., Mencia, R., Garcia, L., Mansilla, N., Alemano, S., Andrade, A. M., Gonzalez, D. H. & Welchen, E. 2019. The mitochondrial oxidation resistance protein AtOXR2 increases plant biomass and tolerance to oxidative stress. Journal of Experimental Botany, 70, 3177–3195.

Danusevicius, D. & Brazaitis, G. 2011. Effect of Provenance on Early Flowering in Norway Spruce. Rural Development in Global Changes, Vol 5, *Book 2*, 5, 30–33.

DE Angeli, A., Zhang, J. B., Meyer, S. & Martinoia, E. 2013. AtALMT9 is a malate-activated vacuolar chloride channel required for stomatal opening in. Nature Communications, 4.

Deng, W. W., Casao, M. C., Wang, P. H., Sato, K., Hayes, P. M., Finnegan, E. J. & Trevaskis, B. 2015. Direct links between the vernalization response and other key traits of cereal crops. Nature Communications, 6.

Deyoung, B. J., Bickle, K. L., Schrage, K. J., Muskett, P., Patel, K. & Clark, S. E. 2006. The CLAVATA1-related BAM1, BAM2 and BAM3 receptor kinase-like proteins are required for meristem function in Arabidopsis. Plant Journal, 45, 1–16.

Dhar, D., Mapa, K., Pudi, R., Srinivasan, P., Bodhinathan, K. & Das, S. 2006. Human ribosomal protein L18a interacts with hepatitis & virus internal ribosome entry site. Archives of Virology, 151, 509–524.

Dutta, A., Choudhary, P., Caruana, J. & Raina, R. 2017. JMJ27, an Arabidopsis H3K9 histone demethylase, modulates defense against *Pseudomonas syringae* and flowering time. Plant Journal, 91, 1015–1028.

Eklöf, H., Bernhardsson, C. & Ingvarsson, P. K. 2026. Contrasting Patterns of Local Adaptation and Adaptive Potential Under Climate Change for Old-Growth and Planted Stands of Norway Spruce (Picea abies). Evolutionary Applications, 19.

Field, D. L., Stankowski, S., Reiter, T., Polechova, J., Bradley, D., Richardson, D. M., Whibley, A., Pal, A., Shipilina, D., Boell, L., Pickup, M., Xue, Y. B., Coen, E. & Barton, N. 2026. Genome-wide cline analysis identifies new locus contributing to a barrier to gene flow across an *Antirrhinum* hybrid zone. Plos Genetics, 22.

Fournier-Level, A., Taylor, M. A., Paril, J. F., Martínez-Berdeja, A., Stitzer, M. C., Cooper, M. D., Roe, J. L., Wilczek, A. M. & Schmitt, J. 2022. Adaptive significance of flowering time variation across natural seasonal environments in. New Phytologist, 234, 719–734.

Gonzalez, J. R., Armengol, L., Sole, X., Guino, E., Mercader, J. M., Estivill, X. & Moreno, V. 2007. SNPassoc: an R package to perform whole genome association studies. Bioinformatics, 23, 644–645.

Grebner, D. L., Bettinger, P., Siry, J. P. & Kevin, B. 2021. Introduction to Forestry and Natural Resources, Elsevier Inc.

Hanano, S. & Goto, K. 2011. TERMINAL FLOWER1 Is Involved in the Regulation of Flowering Time and Inflorescence Development through Transcriptional Repression. Plant Cell, 23, 3172–3184.

Hao, Z. Q., Wang, T. Y., Chen, D. D., Shen, L., Zhang, G. H., Qian, Q. & Zhu, L. 2025. Leucine-Rich Repeat Protein Family Regulates Stress Tolerance and Development in Plants. Rice Science, 32, 32–43.

Hare, P. D., Moller, S. G., Huang, L. F. & Chua, N. H. 2003. LAF3, a novel factor required for normal phytochrome A signaling. Plant Physiology, 133, 1592–1604.

Hord, C. L. H., Chen, C. B., Deyoung, B. J., Clark, S. E. & Ma, H. 2006. The BAM1/BAM2 receptor-like kinases are important regulators of *Arabidopsis* early anther development. Plant Cell, 18, 1667–1680.

Hsieh, W. Y., Liao, J. C., Chang, C. Y., Harrison, T., Boucher, C. & Hsieh, M. H. 2015. The SLOW GROWTH3 Pentatricopeptide Repeat Protein Is Required for the Splicing of Mitochondrial *NADH Dehydrogenase Subunit7* Intron 2 in *Arabidopsis*. Plant Physiology, 168, 490-+.

Huang, M. S., Abel, C., Sohrabi, R., Petri, J., Haupt, I., Cosimano, J., Gershenzon, J. & Tholl, D. 2010. Variation of Herbivore-Induced Volatile Terpenes among Arabidopsis Ecotypes Depends on Allelic Differences and Subcellular Targeting of Two Terpene Synthases, TPS02 and TPS03. Plant Physiology, 153, 1293–1310.

Jamsheer, K. M., Sharma, M., Singh, D., Mannully, C. T., Jindal, S., Shukla, B. N. & Laxmi, A. 2018. FCS-like zinc finger 6 and 10 repress SnRK1 signalling in Arabidopsis. Plant Journal, 94, 232–245.

Jin, J., Tian, F., Yang, D. C., Meng, Y. Q., Kong, L., Luo, J. & Gao, G. 2017. PlantTFDB 4.0: toward a central hub for transcription factors and regulatory interactions in plants. Nucleic Acids Res, 45, D1040–D1045.

Karlgren, A., Gyllenstrand, N., Clapham, D. & Lagercrantz, U. 2013. FLOWERING LOCUS T/TERMINAL FLOWER1-Like Genes Affect Growth Rhythm and Bud Set in Norway Spruce. Plant Physiology, 163, 792–803.

Kertesz, M., Iovino, N., Unnerstall, U., Gaul, U. & Segal, E. 2007. The role of site accessibility in microRNA target recognition. Nature Genetics, 39, 1278–1284.

King, G. J., Chanson, A. H., Mccallum, E. J., Ohme-Takagi, M., Byriel, K., Hill, J. M., Martin, J. L. & Mylne, J. S. 2013. The *Arabidopsis* B3 Domain Protein VERNALIZATION1 (VRN1) Is Involved in Processes Essential for Development, with Structural and Mutational Studies Revealing Its DNA-binding Surface. Journal of Biological Chemistry, 288, 3198–3207.

Landoni, B., SUÁREZ-Montes, P., Habeahan, R. H. F., Brennan, A. C. & Pérez-Barrales, R. 2024. Local climate and vernalization sensitivity predict the latitudinal patterns of flowering onset in the crop wild relative Linum bienne Mill. Annals of Botany, 134, 117–130.

Larsson, A. S., Landberg, K. & Meeks-Wagner, D. R. 1998. The *TERMINAL FLOWER2* (*TFL2*) gene controls the reproductive transition and meristem identity in *Arabidopsis thaliana*. Genetics, 149, 597–605.

Lee, H., Suh, S. S., Park, E., Cho, E., Ahn, J. H., Kim, S. G., Lee, J. S., Kwon, Y. M. & Lee, I. 2000. The AGAMOUS-LIKE 20 MADS domain protein integrates floral inductive pathways in. Genes & Development, 14, 2366–2376.

Li, J. C., Liang, W. J., He, X. Q. & Qian, W. Q. 2025. DNA damage triggers heritable alterations in DNA methylation patterns in Arabidopsis. Molecular Plant, 18, 501–512.

Liu, X., Gao, Y. B., Tang, S. Y., Ben, L. L., Zhang, X. X., Dong, G. C., Zhou, J., Lin, L. S., Yang, Z. F., Zhou, Y., Huang, J. Y. & Yao, Y. L. 2025. A Dual-localized Fructose Bisphosphate Aldolase is Essential for Chloroplast Development and Carbon Metabolism in Rice. Rice, 18.

Luoranen, J., Riikonen, J. & Saksa, T. 2022. Factors affecting winter damage and recovery of newly planted Norway spruce seedlings in boreal forests. Forest Ecology and Management, 503.

Macknight, R., Bancroft, I., Page, T., Lister, C., Schmidt, R., Love, K., Westphal, L., Murphy, G., Sherson, S., Cobbett, C. & Dean, C. 1997. Fca, a gene controlling flowering time in Arabidopsis, encodes a protein containing RNA-binding domains. Cell, 89, 737–745.

Marchler-Bauer, A., Derbyshire, M. K., Gonzales, N. R., Lu, S. N., Chitsaz, F., Geer, L. Y., Geer, R. C., He, J., Gwadz, M., Hurwitz, D. I., Lanczycki, C. J., Lu, F., Marchler, G. H., Song, J. S., Thanki, N., Wang, Z. X., Yamashita, R. A., Zhang, D. C., Zheng, C. J. & Bryant, S. H. 2015. CDD: NCBI’s conserved domain database. Nucleic Acids Research, 43, D222–D226.

Matsui, A., Hashiguchi, K., Suzuki, M. & Zhang-Akiyama, Q. M. 2020. Oxidation resistance 1 functions in the maintenance of cellular survival and genome stability in response to oxidative stress-independent DNA damage. Genes and Environment, 42.

Mei, Q. M., Zheng, Y., Feng, J. Y., Wang, Z. F., Cao, H. L. & Lian, J. Y. 2025. Transcriptome Profiling Revealed Light-Mediated Gene Expression Patterns of Plants in Forest Vertical Structures. Biology-Basel, 14.

Mikola, J. 1982. Bud-set phenology as an indicator of climatic adaptation of Scots pine in Finland. Silvae Fennica, 16, 178–184.

Molmann, J. A., Junttila, O., Johnsen, O. & Olsen, J. E. 2006. Effects of red, far-red and blue light in maintaining growth in latitudinal populations of Norway spruce (*Picea abies*). Plant Cell and Environment, 29, 166–172.

Muthusamy, M., Yoon, E. K., Kim, J. A., Jeong, M. J. & Lee, S. I. 2020. *Brassica Rapa SR45a* Regulates Drought Tolerance via the Alternative Splicing of Target Genes. Genes, 11.

Nikkanen, T. & Ruotsalainen, S. 2000. Variation in flowering abundance and its impact on the genetic diversity of the seed crop in a Norway spruce seed orchard. Silva Fennica, 34, 205–222.

Nilsen, J. 1985. Light climate in Northern areas. Plant Production in the North. Eds. A. Kaurin, O. Juntilla and J Nilsen. Norwegian University Press, Norway, 62–72.

Notivol, E., García-Gil, M. R., Alía, R. & Savolainen, O. 2007. Genetic variation of growth rhythm traits in the limits of a latitudinal cline in Scots pine. Canadian Journal of Forest Research-Revue Canadienne De Recherche Forestiere, 37, 540–551.

Pastore, J. J., Limpuangthip, A., Yamaguchi, N., Wu, M. F., Sang, Y., Han, S. K., Malaspina, L., Chavdaroff, N., Yamaguchi, A. & Wagner, D. 2011. LATE MERISTEM IDENTITY2 acts together with LEAFY to activate. Development, 138, 3189–3198.

Ranade, S. S., Delhomme, N. & García-Gil, M. R. 2019. Transcriptome analysis of shade avoidance and shade tolerance in conifers. Planta, 250, 299–318.

Ranade, S. S. & GARCÍA-Gil, M. R. 2021. Molecular signatures of local adaptation to light in Norway spruce. Planta, 253, 53.

Ranade, S. S. & GARCÍA-Gil, M. R. 2023. Clinal variation in PHY (PAS domain) and CRY (CCT domain)-Signs of local adaptation to light quality in Norway spruce. Plant Cell and Environment, 46, 2391–2400.

Ranade, S. S. & García-Gil, M. R. 2024.Lignin biosynthesis pathway repressors in gymnosperms: differential repressor domains as compared to angiosperms. Forestry Research, 4.

Ranade, S. S. & GARCÍA-Gil, M. R. 2025. Metabolomic profiling of shade response and in silico analysis of PAL homologs imply the potential presence of bifunctional ammonia lyases in conifers. Physiologia Plantarum, 177.

Ranade, S. S., Seipel, G., Gorzsás, A. & García-Gil, M. R. 2022. Enhanced lignin synthesis and ecotypic variation in defense-related gene expression in response to shade in Norway spruce. Plant Cell and Environment, doi: 10.1111/pce.14387.

Sharma, A., Gupta, A., Ramakrishnan, M., Van Ha, C., Zheng, B. S., Bhardwaj, M. & Tran, L. S. P. 2023. Roles of abscisic acid and auxin in plants during drought: A molecular point of view. Plant Physiology and Biochemistry, 204.

Skroppa, T., Tollefsrud, M. M., Sperisen, C. & Johnsen, O. 2010. Rapid change in adaptive performance from one generation to the next in Picea abies-Central European trees in a Nordic environment. Tree Genetics & Genomes, 6, 93–99.

Sogaard, G., Johnsen, O., Nilsen, J. & Junttila, O. 2008. Climatic control of bud burst in young seedlings of nine provenances of Norway spruce. Tree Physiology, 28, 311–320.

Ståldal, V., Cierlik, I., Chen, S., Landberg, K., Baylis, T., Myrenås, M., Sundström, J. F., Eklund, D. M., Ljung, K. & Sundberg, E. 2012. The Arabidopsis thaliana transcriptional activator STYLISH1 regulates genes affecting stamen development, cell expansion and timing of flowering. Plant Molecular Biology, 78, 545–559.

Sundell, D., Mannapperuma, C., Netotea, S., Delhomme, N., Lin, Y. C., Sjodin, A., VAN DE Peer, Y., Jansson, S., Hvidsten, T. R. & Street, N. R. 2015. The Plant Genome Integrative Explorer Resource: PlantGenIE.org. New Phytologist, 208, 1149–56.

Svystun, T., Lundströmer, J., Berlin, M., Westin, J. & Jönsson, A. M. 2021. Model analysis of temperature impact on the Norway spruce provenance specific bud burst and associated risk of frost damage. Forest Ecology and Management, 493.

Tao, Y., Wu, L. L., Volodymyr, V., Hu, P., Hu, H. Y. & Li, C. W. 2024. Identification of the ribosomal protein L18 (RPL18) gene family reveals that TaRPL18-1 positively regulates powdery mildew resistance in wheat. International Journal of Biological Macromolecules, 280.

Wang, X., Bernhardsson, C. & Ingvarsson, P. K. 2020. Demography and Natural Selection Have Shaped Genetic Variation in the Widely Distributed Conifer Norway Spruce (Picea abies). Genome Biology and Evolution, 12, 3803–3817.

Westin, J., Sundblad, L. G., Strand, M. & Hällgren, J. E. 2000a. Phenotypic differences between natural and selected populations of *Picea abies*. I. Frost hardiness. Scandinavian Journal of Forest Research, 15, 489–499.

Westin, J., Sundblad, L. G., Strand, M. & Hällgren, J. E. 2000b. Phenotypic differences between natural and selected populations of *Picea abies*. II. Apical mitotic activity and growth related parameters. Scandinavian Journal of Forest Research, 15, 500–509.

Williams, C. G. 2009. Introducing Conifers. Conifer Reproductive Biology. Dordrecht: Springer.

Wu, X. Y., Xia, M., Su, P., Zhang, Y. F., Tu, L. C., Zhao, H., Gao, W., Huang, L. Q. & Hu, Y. T. 2024. MYB transcription factors in plants: A comprehensive review of their discovery, structure, classification, functional diversity and regulatory mechanism. International Journal of Biological Macromolecules, 282.

Wu, Y., Li, X., Zhang, J. N., Zhao, H. Q., Tan, S. L., Xu, W. H., Pan, J. Q., Yang, F. & Pi, E. X. 2022. ERF subfamily transcription factors and their function in plant responses to abiotic stresses. Frontiers in Plant Science, 13.

Xu, J. J., Lei, Y., Zhang, X. F., Li, J. X., Lin, Q. P., Wu, X. D., Jiang, Y. G., Zhang, W. Y., Qian, R. T., Xiong, S. Y., Tan, K., Jia, Y., Zhou, Q., Jiang, Y., Fan, H., Huang, Y. B., Wang, L. J., Liu, J. Y., Kong, Y., Zhao, Q., Yang, L., Liu, J. X., Hu, Y. H., Zhan, S., Gao, C. X. & Chen, X. Y. 2025. Design of CoǪ10 crops based on evolutionary history. Cell, 188, 1941–1954.

Yant, L., Mathieu, J., Dinh, T. T., Ott, F., Lanz, C., Wollmann, H., Chen, X. M. & Schmid, M. 2010. Orchestration of the floral transition and Floral development in *Arabidopsis* by the bifunctional transcription factor APETALA2. Plant Cell, 22, 2156–2170.

Yoo, S. Y., Kim, Y., Kim, S. Y., Lee, J. S. & Ahn, J. H. 2007. Control of Flowering Time and Cold Response by a NAC-Domain Protein in. Plos One, 2.

Zhao, G. C., Shi, J. X., Liang, W. Q. & Zhang, D. B. 2016. ATP binding cassette G transporters and plant male reproduction. Plant Signaling & Behavior, 11.

Zhao, T. Y., Yang, X., Yang, X. Y., Rao, P., An, X. M. & Chen, Z. 2021. Identification of key flowering-related genes and their seasonal expression in reproductive buds suggests dual roles in floral development and dormancy. Industrial Crops and Products, 161.

Zuo, Z. F., Lee, H. Y. & Kang, H. G. 2023. Basic Helix-Loop-Helix Transcription Factors: Regulators for Plant Growth Development and Abiotic Stress Responses. International Journal of Molecular Sciences, 24.

