## Supplementary Figures S1-S25 for "Genome-wide latitudinal variation associated with defence and reproductive pathways linked to Norway spruce adaptation"

**Figures S1-S20, S25, S26 represent alignments of the Norway spruce genes with the model plant *Arabidopsis*; CLUSTAL O(1.2.4) multiple sequence alignment was used for all alignments. SNP highlighted in green represent missense and those highlighted in yellow represent synonymous mutations, unless specified otherwise.**

**Figures S22-S24 represent Gene Ontology (GO) enrichment analysis**

**Figure S1** Alignment between Norway spruce 60S ribosomal protein L18 (RPL18C) (MA_3674g0020) and *Arabidopsis thaliana* RPL18C (AT5G27850). Norway spruce RPL18C is spread across two contigs, refer Figure S2.

```
MA_3674g0020      MGIDLIAGGKSKQTKRTAPKSEDIYLKLLVKLYRFLVRRTSSRFNAASAVAPRTDSLLLT  60
AT5G27850         MGIDLIAGGKSKKTKRTAPKSDDVYLKLLVKLYRFLVRRSNSNFNAVILKRLFMSKVN--  58
                  ************:********:*:***************:.*.***.       ..:

MA_3674g0020      STRCFFLVFCVVSLMPFLLVYND--------------------------SYERRP-----  89
AT5G27850         --KA---PLSLSRLVEFMTGKDDKIAVLVGTITDDLRVHEIPAMKVTALRFTERARARIE  113
                    :.    :.:  *: *:   :*                           : .*

MA_3674g0020      ------------------------------------------------------------  89
AT5G27850         KAGGECLTFDQLALRAPLGQNTVLLRGPKNSREAVKHFGPAPGVPHSNTKPYVRHKGRKF  173


MA_3674g0020      -------------- 89
AT5G27850         EKARGKRKSRGFKV 187
```

L MA_3674_19426, cline 0.6
C > T
CTC > TTC
Leu(L) > Phe(F)
Leu74Phe
Non-polar hydrophobic > Non-polar hydrophobic

**Figure S2** Alignment between Norway spruce RPL18C (MA_5630845g0010) and *Arabidopsis thaliana* RPL18C (AT5G27850). Norway spruce RPL18C is spread across two contigs, refer Figure S1.

```
MA_5630845g0010      ------------------------------------------------------------     0
AT5G27850            MGIDLIAGGKSKKTKRTAPKSDDVYLKLLVKLYRFLVRRSNSNFNAVILKRLFMSKVNKA     60


MA_5630845g0010      --------------DDKIAVVVGTITDDNRVYELPPVKVTALRFTETARARILKAGGECL     46
AT5G27850            PLSLSRLVEFMTGKDDKIAVLVGTITDDLRVHEIPAMKVTALRFTERARARIEKAGGECL     120
                                   ******:******* **:*:* :********* ***** *******

MA_5630845g0010      TFDQLALRAPLGQNTVLLRGPKSHREAVKHFGRAPGVPHSHTKPYVRSKGRKYEMARGRR     106
AT5G27850            TFDQLALRAPLGQNTVLLRGPKNSREAVKHFGPAPGVPHSNTKPYVRHKGRKFEKARGKR     180
                     **********************. ******** *******:****** ****:* ***:*

MA_5630845g0010      NSRGFKV     113
AT5G27850            KSRGFKV     187
                     :******
```

**Figure S3** Alignment between Norway spruce Long after far-red 3 (LAF3) (MA_9446113g0010) and *Arabidopsis thaliana* LAF3 (AT3G55850). Norway spruce LAF3 is spread across two contigs, refer Figure S4.

```
MA_9446113g0010      ------------------------------------------------------------      0
AT3G55850            ADSMAIRNGRILKVGSFATLKGFIGDGTMEVNLEGKIVVPGLIDSHVHLISGGLQMAQVG      120


MA_9446113g0010      ------------------------------------------------------------      0
AT3G55850            LRGVSQKDEFCKMVKDAVQNAKEGSWILGGGWNNDFWGGELPSASWIDEISPRNPVWLIR      180


MA_9446113g0010      ------------------------------------------------------------      0
AT3G55850            MDGHMALANSLALKIAGVISLTEDPVGGTIMRMPSGEPTGLLIDAAMELVTPWVKEISVD      240


MA_9446113g0010      ------------------------------------------------------------      0
AT3G55850            ERREALFRASKYALTRGVTTVIDLGRYFPGTTDELSWKDFQDVYLYADSSKKMMIRTCLF      300


MA_9446113g0010      ------------------------------------------------------------      0
AT3G55850            FPITTWSRLLDLKLQKGSVLSEWLYLGGVKAFIDGSLGSNSALFYEEYIDTPNNYGLEVM      360


MA_9446113g0010      ------------------------------------------------------------      0
AT3G55850            DPEKLSNFTMAADKSGLQVAIHAIGDKANDMILDMYESVAAANGDRDRRFRIEHAQHLAP      420


MA_9446113g0010      -------------MIYPAHLLDDAISATKKLGEKRALQESYLFGSLLLSNTRLAFGSDWP      47
AT3G55850            GSANRFGQLHIVASVQPDHLLDDADSVAKKLGSERAVKESYLFQSLLNGNALLALGSDWP      480
                                   : * ****** *.:****.:**::***** *** .*: **:*****

MA_9446113g0010      VADLNPLGGIKAATERMRSNWQRPWIESERITVDAALN----------------------      85
AT3G55850            VADINPLHSIRTAVKRIPPKWDHAWIPSERISFTDALIAQTISAARAAFLDHHLGSLSPG      540
                     ***:*** .*::*.:*:  :*:: ** ****:.  **

MA_9446113g0010      ------------------------------------   85
AT3G55850            KLADFVILSTNSWDEFSKDVSASVLATYVGGKQLYP   576
```

L MA_9446113_1550, cline 0.6
T > C
TTA > TCA
Leu(L) > Ser(S)
Leu35Ser
Non-polar hydrophobic > Polar hydrophilic

**Figure S4** Alignment between Norway spruce LAF3 (MA_957354g0010) and *Arabidopsis thaliana* LAF3 (AT3G55850). Norway spruce LAF3 is spread across two contigs, refer Figure S3.

```
MA_957354g0010      ------------------------------------------------------------      0
AT3G55850           MNLFVIVSAAVFLLISVAYLPLLNDLYWSTLKSLTPPAGIVADLLVTNGTIFTSDSSLPF      60


MA_957354g0010      ----------------------------------------MLLATSVYHYTPHDNVKQPS      20
AT3G55850           ADSMAIRNGRILKVGSFATLKGFIGDGTMEVNLEGKIVVPGLIDSH-VHLIS-GGLQMAQ      118
                                                             *: :   *    ..::  .

MA_957354g0010      TGAFFAQKEVSKCR---YILRGMERGHWVLGGGWNNDNWGGELPQASWIDNITHDHPVWL      77
AT3G55850           VGLRGVSQKDEFCKMVKDAVQNAKEGSWILGGGWNNDFWGGELPSASWIDEISPRNPVWL      178
                    .*   ..:: . *:     ::. :.* *:******** ******.*****:*:  :****

MA_957354g0010      SRMDGHMGLANSLALAKAGIYGGSRDIVGGAIIKSNDGAPTGLLVDAAMQLLLPHIPEAT      137
AT3G55850           IRMDGHMALANSLALKIAGVISLTEDPVGGTIMRMPSGEPTGLLIDAAMELVTPWVKEIS      238
                     ******.*******  **: . :.* ***:*::  .* *****:****:*: * : * :

MA_957354g0010      VQKRRDAMIRASKLGLSRGVTAVVDFGRYFPGSSTRHVWDDFSEVYKWADETGHMLLRVL      197
AT3G55850           VDERREALFRASKYALTRGVTTVIDLGRYFPGTTDELSWKDFQDVYLYADSSKKMMIRTC      298
                    *::**:*::**** .*:****:*:*:******:: .  *.**.:** :**.: :*::*.

MA_957354g0010      REAGQ---------------GL------EEVHAFLEEDLLKTTLEEEVHIFLEEAH--IN      234
AT3G55850           LFFPITTWSRLLDLKLQKGSVLSEWLYLGGVKAFIDG-----SLGSNSALFYEEYIDTPN      353
                                         *        *:**::      :* .:  :* **     *

MA_957354g0010      SWAQVVQGRDKLPPPPS---CCG-------------------------------------      254
AT3G55850           NYGLEVMDPEKLSNFTMAADKSGLQVAIHAIGDKANDMILDMYESVAAANGDRDRRFRIE      413
                    .:.  * . :**         .*

MA_957354g0010      ------------------------------------------------------------      254
AT3G55850           HAQHLAPGSANRFGQLHIVASVQPDHLLDDADSVAKKLGSERAVKESYLFQSLLNGNALL      473


MA_957354g0010      ------------------------------------------------------------      254
AT3G55850           ALGSDWPVADINPLHSIRTAVKRIPPKWDHAWIPSERISFTDALIAQTISAARAAFLDHH      533


MA_957354g0010      -------------------------------------------    254
AT3G55850           LGSLSPGKLADFVILSTNSWDEFSKDVSASVLATYVGGKQLYP    576
```

S MA_957354_5489, cline 0.4
C > T
TCA > TTA
Ser(S) > Leu(L)
Ser31Leu
Polar hydrophilic > Non-polar hydrophobic

**Figure S5** Alignment between Norway spruce ZINC 4 FINGER DNA 3'-PHOSPHOESTERASE (ZDP) (MA_39122g0010) and *Arabidopsis thaliana* ZDP (AT3G14890). HAD_PNP domain is showed in **bold**.

```
MA_39122g0010      ------------------------------------------------------------ 0
AT3G14890          MITVAPFVSLRFQFPLYIINRSTLFFRERTQYLVYNTCHILRTTAKTMPVVAEYAKSNRS 60


MA_39122g0010      ------------------------------------------------------------ 0
AT3G14890          SCRSCSNKIAVKSLRLGLISKGRGGVDMTRWHHFDCFPTDSESIASVDDIQGLSALEKED 120


MA_39122g0010      ------------------------------------------------------------ 0
AT3G14890          QDALTKLVEQCGKVPAKKPDEKKGKAKKHIMGPKGLTKAATSSKVIADNAKSSRSSCNRC 180


MA_39122g0010      ------------------------------------------------------------ 0
AT3G14890          SQTIVSKDLRVGLVTEDSRGFDITRWHHLGCFPIDFHPIDSVEDIGGYSSLEKGDQMELK 240


MA_39122g0010      ------------------------------------------------------------ 0
AT3G14890          YLAEVNKKDKTLIDDVQKMDEGDDEAIADNELTEETKKGKHSPVAKLVEQPGEPAKEDED 300


MA_39122g0010      ------------------------------------------------------------ 0
AT3G14890          EESKKPASDEISEQKTKDVKNSPDSSKVISEYAKSSRSTCKKCSQTIAAKELRLGLVTRN 360


MA_39122g0010      ------------------------------------------------------------ 0
AT3G14890          FRGFDMKQWHHLGCFPVDSDPIVSVEDIGGFSELQSGDQDALKELVQQCGKQTLVDKMDE 420


MA_39122g0010      ------------------------------------------------------------ 0
AT3G14890          DNDDTEAKIKLTEETNKRKHSEVGEMVEEDESLTKAKQQMAKTHKVNMSESTSQVEVEAE 480


MA_39122g0010      ------------------------------------------------------------ 0
AT3G14890          ITLSASDVKDKYRDANLLPKWKAFETVIFLERDDGLNDSEKIAAFDFDGCLAKTSVKIVG 540


MA_39122g0010      ------------------------------------------------------------ 0
AT3G14890          ADAWSLMYPSIPEKLQSLHDQGYKLVIFTNESNIDRWKNKRQAAVDSKIGRLNSFIERVK 600


MA_39122g0010      ---------------MIEGTGDAFRKPKSGMWRLMEPHFNSEIIINMEQSFCVGDAAGRI 45
AT3G14890          VPIQVFIACGVSSSGGKGGKDDLYRKPKAGMWQLMKKHFNSGIAIDMDKSFYVGDAAGRK 660
                                     *..* :****:***:**: **** * *:*::** *******

MA_39122g0010      NDHSDADICFAKDVGLKFPSSDGCLQI------- 72
AT3G14890          MDHSDADIKFAQASGLKFFTPEEYFIPSSTSPGT 694
                    ******* **:  **** : :  :
```

P MA_39122_15150, cline 0.5
C > G
CCA > CGA
Pro(P) > Arg(R)
Pro22Arg
Non-polar hydrophobic > Polar hydrophilic

N MA_39122_15124, cline 0.5
A > G
AAC > GAC
Asn(N) > Asp(D)
Asn31Asp
Polar hydrophilic > Polar hydrophilic

C MA_39122_15009, cline 0.5
G > A
TGT > TAT
Cys(C) > Tyr(Y)
Cys69Tyr
Polar hydrophilic > Polar hydrophilic

**Figure S6** Alignment between Norway spruce Coenzyme Q9 (COQ9) (MA_9293920g0010) and *Arabidopsis thaliana* COQ9 (AT1G19140). Norway spruce COQ9 is spread across two contigs, see Figure S7 for complete gene.

```
MA_9293920g0010      ------------------------------------------------------------      0
AT1G19140            MYRTAAKRLLAGGGITTTRLLRLPKLRSTIIPSSYTSGLCTSSIGGNTESPMGKQSVNPN      60


MA_9293920g0010      ------------------------------------------------------------      0
AT1G19140            QSGPTASSSTGTGEGQRRHESRKPRAEFQEEQARVLSASLRHVPRLGWTEEAMMAGSRDV      120


MA_9293920g0010      ---------------------MDDCLQRLLGEIESREEELSNLVFHDRIAKIIRIRLEMQ      39
AT1G19140            GVSPSIVGSFSRKEAALVEFFMDECLQLLMDRIDSG-LDLQNLIPSERISKLIRIRLEMQ      179
                                          **:*** *:..*:*   :*.**:  :**:*:********

MA_9293920g0010      VPYISKWPQALSIQANPLNLPTSFKQRAVLVDEIWHASGDRSTDLDWYAKRTLLGGIYAA      99
AT1G19140            VPYMSKWPQALSIQAHPLNVPTSFKQRAMLVDEIWHAVGDGASDLDWYVKRTILGGVYST      239
                     ***:***********:***:********:******** ** ::*****.***:***:*::

MA_9293920g0010      TELYMLTDYSPEFRNTWIFLDRRVSDAVDCRKTAQEAAQLATAIGAGIGNTVQAFLRRQT      159
AT1G19140            TEIYMLTDDSLEHRDTWAFLDDRVKDAFDLKKSIQEAKYFAEDIGAGVGKSVQGLMNGVM      299
                     **:***** * *.*:** *** **.**.* :*: ***   :*   ****:*::**.::.

MA_9293920g0010      Q----------- 160
AT1G19140            QTMSRRSGGSAF 311
                     *
```

S MA_9293920_3163, cline 0.5
T > G
TCT > GCT
Ser(S) > Ala(A)
Ser77Ala
Polar hydrophilic > Non-polar hydrophobic

**Figure S7** Alignment between Norway spruce Coenzyme Q9 (COQ9) (MA_113286g0010) and *Arabidopsis thaliana* COQ9 (AT1G19140). Norway spruce COQ9 is spread across two contigs, see Figure S6 for complete gene.

```
MA_113286g0010      ---MNIIRWSFANGIRRTRWLSISSPIHRDLNNPLLFHPSQSPLPSNNRPLPYNL----R       53
AT1G19140           MYRTAAKRLLAGGGITTTRLLRLPKL--R----------------STIIPSSYTSGLCTS       42
                           *   ..**  ** * : .   *                *.  *  *.

MA_113286g0010      ALSDNTSSSSIHEDVKPE---P----TQTVSQEQQTEEEKKTNHDIKDDQTRVLQAALEH       106
AT1G19140           SIGGNTESPMGKQSVNPNQSGPTASSSTGTGEGQRRHESRKPRAEFQEEQARVLSASLRH       102
                    ::..**.*   ::.*:*:   *    :  ..: *: .*.:* .  ::::*:***.*:*.*

MA_113286g0010      VPTLGWTEAAMVAGAREVGLSPSIVGAFPRKEAALVEVFIVPLSF---------------       151
AT1G19140           VPRLGWTEEAMMAGSRDVGVSPSIVGSFSRKEAALVEFFMDECLQLLMDRIDSGLDLQNL       162
                    ** ***** **:**:*:**:******:* ********.*:

MA_113286g0010      ------------------------------------------------------------       151
AT1G19140           IPSERISKLIRIRLEMQVPYMSKWPQALSIQAHPLNVPTSFKQRAMLVDEIWHAVGDGAS       222


MA_113286g0010      ------------------------------------------------------------       151
AT1G19140           DLDWYVKRTILGGVYSTTEIYMLTDDSLEHRDTWAFLDDRVKDAFDLKKSIQEAKYFAED       282


MA_113286g0010      -----------------------------    151
AT1G19140           IGAGVGKSVQGLMNGVMQTMSRRSGGSAF    311
```

L MA_113286_17254, cline 0.1
C > T
CTT > TTT
Leu(L) > Phe(F)
Leu33Phe
Non-polar hydrophobic > Non-polar hydrophobic

P MA_113286_17276, cline 0.1
C > T
CCT > CTT
Pro(P) > Leu(L)
Pro40Leu
Non-polar hydrophobic > Non-polar hydrophobic

S MA_113286_17342, cline 0.1
C > A
TCT > TAT
Ser(S) > Tyr(Y)
Ser62Tyr
Polar hydrophilic > Polar hydrophilic

K MA_113286_17413, cline 0.1
A > G
AAG > GAG
Lys(K) > Glu(E)
Lys86Glu
Polar hydrophilic > Polar hydrophilic

**Figure S8** Alignment between Norway spruce SERINE/ARGININE RICH-LIKE PROTEIN 45A (SR45a) (MA_42236g0030) and *Arabidopsis thaliana* SR45a (AT1G07350).

```
MA_42236g0030      MLVLAIMLVKMGGPSYTAGFFVFEIAAEENWLRSLQKSGMALESPTMSWEKKENLPRESK 60
AT1G07350          ------------------------------------------------------------ 0


MA_42236g0030      EDHGRRSANLDLNRADEGDEYDYHPYKEEPEKTEGNDNQQNRAFSPRRDQAGDYTGRIYD 120
AT1G07350          ------------------------------------------------------------ 0


MA_42236g0030      SPSKSSPGQSPRLARDRSPSPVRVKSPSPVKYISTSQSRHISSSPTNYTSNLSDSHQISS 180
AT1G07350          ------------------------------------------------------------ 0


MA_42236g0030      PAKPKSPLSINHRSPSPTKHSSPSSGKLSSASPAKYTSPSPERRRSSPPQRDVSSSLKRH 240
AT1G07350          ------------------------------------------------------------ 0


MA_42236g0030      RSSPRRDVAPSPGRHRSSPPRDLPHSPGTHMSSSPRREISPSPGRHRSSPPRRDVSPSPG 300
AT1G07350          ------------------------------------------------------------ 0


MA_42236g0030      RRRSSPPRRDVSPSSGSYGSSPPWRELSASPGRHRTTPSKRRSPVGRYISPSPKRRRYDR 360
AT1G07350          ------------------------------MGK----REIHFTPVGRQVQRV-------- 18
                                                  *:     . : :**** :.

MA_42236g0030      SRTPTGRRGVSPGYGRRGYRDRSQSRSPPRDYYRRSPRGKYSPRSRYSPRRRTPPRFQSR 420
AT1G07350          ----------------LEYPLRLENR----------SPMSYSRRSRYSPSLSP---YDKR 49
                                     *  * :.*             .** ******       ::.*

MA_42236g0030      RR--SPRRRQWSPPPNRNTGIGKPGNNLFVAGFSFDTSERDLERKFSKFGRVTDVRVVRD 478
AT1G07350          RGRSVSRSLSRSPTRSVSSDAENPGNSLYVTGLSHRVTERDLEDHFAKEGKVTDVHLVLD 109
                   *     *  . **  . .:.  :***.*:*:*:*. .:***** :*:* *:****::* *

MA_42236g0030      KRSGDSRGFGFLTLERDEDADAAIRRRHQLDTEVDSFVCIISPLLMFGILKQIQPKIWLS 538
AT1G07350          PWTRESRGFGFISMKSVGDANRCIRSLDHSVLQGR-VITVEK--ARRRRGRTPTPGKYLG 166
                     : :******::::   **: .**  .:   :   .: : .        :   *  :*.

MA_42236g0030      FGCTRSSHDTNYFLKVAQRLSVPFSICS-------------------------------- 566
AT1G07350          LRTARGRHKSPSYSP---RR---SVSCSRSRSRSYSSDRGRSYSPSYGRRGRSSSYSPFY 220
                   :  :*. *.:  :     *       **

MA_42236g0030      ------------------------------------------------------------ 566
AT1G07350          RRRRFYSPSRSPSPDDRYNRRRDRSYSPYYRRRDRSRSYSRNCRARDRSPYYMRRYRSRS 280


MA_42236g0030      ------------------------------------------------------------ 566
AT1G07350          RSYSPRYRARDRSCSPYYRGRDRSYSPHYQGRDRSYSPESRYYRRHRSVSGSVSPGGRSM 340


MA_42236g0030      ------------------------------------------      566
AT1G07350          SRSISPRKGRKESRSKSRRHDRQSSMCHSRSARSSTSRSVSP      382
```

S MA_42236_32143, cline 0.5
G > T
AGC > ATC
Ser(S) > Ile(I)
Ser209Ile
Polar hydrophilic > Non-polar hydrophobic

T MA_42236_32119, cline 0.1
C > T
ACT > ATT
Thr(T) > Ile(I)
Thr217Ile
Polar hydrophilic > Non-polar hydrophobic

**Figure S9** Alignment between Norway spruce ATP-BINDING CASSETTE G26 (ABCG26) (MA_16472g0020) and *Arabidopsis thaliana* ABCG26 (AT3G13220).

```
MA_16472g0020      MSGVRLRCLLDDEPRDSKAAELRFFGSRKSLDGERGRDEGPSVQIPEPSSTFTHVPDIPP 60
AT3G13220          ------------------------MEIRRST---EEVEENHVMQITGSNGIVHNMEFMPQ 33
                                           :  *:*    .  :*.  :**   .. . ::  :*

MA_16472g0020      ------FRSSSKAHRHSYDIETANSSPLPIFIKFEDLAYTVQYNRPPGTWNPLSQLR--H 112
AT3G13220          AYLRNQYSSEIDIDEEFVSTYPLEDAPLPIFLKFEDVEYKVRNSHASSANLVKTMVSKVV 93
                         : *. . ...  .    :.:*****:****: *.*: .:  .:    : :

MA_16472g0020      MGGSQNQSMSKSILNGITGSVAPGQILALMGPSGSGKTTLLKIVGGRIHGDIKGSITYND 172
AT3G13220          THTNPDPDGYKHILKGITGSTGPGEILALMGPSGSGKTTLLKIMGGRLTDNVKGKLTYND 153
                      . : .  * **:*****..**:******************:***: .::**.:****

MA_16472g0020      IPYSTALKRRYASISN--IFLCGFAHLEICA----------------------------- 201
AT3G13220          IPYSPSVKRRIGFVTQDDVLLPQLTVEETLAFAAFLRLPSSMSKEQKYAKIEMIIKELGL 213
                   **** ::*** . :::  ::*  ::  *  *

MA_16472g0020      ------------------------------------------------------------ 201
AT3G13220          ERCRRTRVGGGFVKGISGGERKRASIAYEILVDPSLLLLDEPTSGLDSTSATKLLHILQG 273


MA_16472g0020      ------------------------------------------------------------ 201
AT3G13220          VAKAGRTVITTIHQPSSRMFHMFDKLLLISEGHPAFYGKARESMEYFSSLRILPEIAMNP 333


MA_16472g0020      ------------------------------------------------------------ 201
AT3G13220          AEFLLDLATGQVSDISLPDELLAAKTAQPDSEEVLLKYLKQRYKTDLEPKEKEENHRNRK 393


MA_16472g0020      ------------------------------------------------------------ 201
AT3G13220          APEHLQIAIQVKKDWTLSWWDQFLILSRRTFRERRRDYFDKLRLVQSLGVAVVLGLLWWK 453


MA_16472g0020      ------------------------------------------------------------ 201
AT3G13220          SKTDTEAHLRDQVGLMFYICIFWTSSSLFGAVYVFPFEKIYLVKERKAEMYRLSVYYVCS 513


MA_16472g0020      ------------------------------------------------------------ 201
AT3G13220          TLCDMVAHVLYPTFFMIIVYFMAEFNRNIPCFLFTVLTILLIAITSQGAGEFLGASVLSI 573


MA_16472g0020      ------------------------------------------------------------ 201
AT3G13220          KRAGMIASLVLMLFLLTGGYYVQHIPKFMQWLKYLSFMHYGFRLLLKVQYSADQLFECGS 633


MA_16472g0020      ----------------------------------------------------  201
AT3G13220          KGGCRTLQSSSSFDTINLNGGLQELWVLLAMAFGYRLCAYFCLRKKISICHL  685
```

E MA_16472_28527, cline 0.5

A > T
GAA > GAT
Glu(E) > Asp(D)
Glu34Asp
Polar hydrophilic > Polar hydrophilic

**Figure S10** Alignment between Norway spruce OXIDATION RESISTANCE 1 (OXR1) (MA_697287g0010) and *Arabidopsis thaliana* OXR1 (AT4G39870).

```
MA_697287g0010      -------------------------------------------MEKKRRHQKTFSEKGSS      17
AT4G39870           MGKHKSFRSKAVHFVTDLTAGLLNPISDKPSSAHPPPPLPDEEDESKRNQLESTTAEQPK      60
                                                                *.**.: :: : :  .

MA_697287g0010      DQAGGPDTTSFMSFIYSFLTCSEASVRRAEDGEKDECNIEMNEQYSTK-----GSVGSRK      72
AT4G39870           DLVDEPDTSSFSAFLGSLLSSDPKDKRKDQDPEDEEDEEEDEEEDSEAETSDTSSSSANP      120
                    * .. ***:** :*: *:*:..  . *: :* *.:* : * :*: *       .* .:.

MA_697287g0010      RQSAKSK----------CSSGKKH----YQGRNSPQSSNQEDAHGAGNGDEGDWHFLTDH      118
AT4G39870           TRTMKETTSGGAAKKSFLSKYKQHFRNFYQAVKFPGVKERKGNSDV-IPDDEETEYYDGL      179
                     :: *..           *. *:*    **. : *  .:::.  ..   *: : .:  .

MA_697287g0010      DIK-----SEKGSHKSIQQDMLLEMSEPSLLLSKHFSSYLCSELPTLVQGRKWVLLYSTW      173
AT4G39870           EMKPMQNNNVKEEVTVVVQAIIPEISEPSLLLSEQSRRSLYTSLPALVQGRKWILLYSTW      239
                    ::*     . * . . : * :: *:********::    * :.**:*******:******

MA_697287g0010      KHGISLHTLYRRSTLSPGPCLL--------------------------------------      195
AT4G39870           RHGISLSTLYRKSLLWPGLSLLVVGDRKGSVFGGLVEAPLIPTDKKYQGTNSTFVFTNKS      299
                    :***** ****:* * ** .**

MA_697287g0010      ------------------------------------------------------------      195
AT4G39870           GQPTIYRPTGANRFYTLCSKEFLALGGGGRFALYLDSELLSGSSAYSETYGNSCLADSQD      359


MA_697287g0010      -----------------------------------     195
AT4G39870           FDVKEVELWGFVYGSKYDEILAHSKTMEPGLCRWS     394
```

R MA_697287_2691, cline 0.4
G > A
AGG > AAG
Arg(R) > Lys(K)
Arg71Lys
Polar hydrophilic > Polar hydrophilic

S MA_697287_2518, cline 0.5
T > G
TCA > GCA
Ser(S) > Ala(A)
Ser129Ala
Polar hydrophilic > Non-polar hydrophobic

**Figure S11** Alignment between Norway spruce ALUMINUM-ACTIVATED MALATE TRANSPORTER 9 (ALMT9) (MA_10243484g0010) and *Arabidopsis thaliana* ALMT9 (AT3G18440).

```
MA_10243484g0010      ------------------------------------------------------------    0
AT3G18440             MAAKQGSFRHGILEKRERLLSNNGFSDFRFTDIESNDLLENENCGRRTRLCCCCSCGNLS    60


MA_10243484g0010      ------------------------------------------------------------    0
AT3G18440             EKISGVYDDAKDVARKAWEMGVSDPRKIVFSAKIGLALTIVALLIFYQEPNPDLSRYSVW    120


MA_10243484g0010      ------------------------------------------------------------    0
AT3G18440             AILTVVVVFEFTIGATLSKGFNRALGTLSAGGLALGMAELSTLFGDWEEIFCTLSIFCIG    180


MA_10243484g0010      ------------------------------------------------------------    0
AT3G18440             FLATFMKLYPSMKAYEYGFRVFLLTYCYILISGFRTGQFIEVAISRFLLIALGAGVSLGV    240


MA_10243484g0010      ------------------------------------------------------------    0
AT3G18440             NMFIYPIWAGEDLHNLVVKNFMNVATSLEGCVNGYLRCLEYERIPSKILTYQASEDPVYK    300


MA_10243484g0010      ------------------------------------------------------------    0
AT3G18440             GYRSAVESTSQEESLMSFAIWEPPHGPYKSFNYPWKNYVKLSGALKHCAFTVMALHGCIL    360


MA_10243484g0010      ------------------MNIVAEASNVILELADSIKTMTRITRINLMTHRLKRTVEELQN    43
AT3G18440             SEIQAPEERRQVFRQELQRVGVEGAKLLRELGEKVKKMEKLGPVDLLFE-VHLAAEELQH    419
                                        .: .*.:::: **.:.:*.* ::  ::*: . :: :.****:

MA_10243484g0010      CFKSEPGLFMDNKRWQIVQQTPNP----KIINFKITPPSESINPQV----------SRIE    89
AT3G18440             KIDKKSYLLVNSECWEIGNRATKESEPQELLSLEDSDPPENHAPPIYAFKSLSEAVLEIP    479
                       :..:  *:::.: *:* ::: :     :::.:: : * *.  * :           .*

MA_10243484g0010      KADGEANQK-----PIISSKGKIWPRTDIPVTSDDSTVGRTHQVQNHETIPFMDTLSIAT    144
AT3G18440             PSWGEKNHREALNHRPTFSKQVSWPARLVLPPHLETTNGASPLVETTKTYESASALSLAT    539
                       : ** *::         **   **   :     ::* * :  *:. :*    .:**:**

MA_10243484g0010      IACLLTKVVARLETAIIAVDDLGELANFTSIID--DK--PIFR----------------    183
AT3G18440             FASLLIEFVARLQNVVDAFKELSQKANFKEPEIVTTGTDVEFSGERVGLGQKIRRCFGM    598
                      :*.** :.****:..: *..:*.: ***..           *
```

V MA_10243484_1744, cline 0.5

G > T

GTA > TTA

Val(V) > Leu(L)

Val122Leu

Non-polar hydrophobic > Non-polar hydrophobic

**Figure S12** Alignment between Norway spruce TERPENE SYNTHASE 02 (TPS02) (MA_10426551g0010) and *Arabidopsis thaliana* TPS02 (AT4G16730).

```
MA_10426551g0010      MINSVELAEVMAQISKSLSRPANWNKESSIVSDIHGNMRDDGWIQSLSSPYGAPAYRERA    60
AT4G16730             -----------------------------MGSSISPLAREYIYVKEVETAEKAILFKEEV    31
                                                   : *.*    *:  :::.:.:   *  ::*..

MA_10426551g0010      EKLIEEIKHLLFRYMEDRCNDDLIKRAQIVDMVQCLGIDRHFQHELRGALDCLYRCWNER    120
AT4G16730             RKTLNE-------------IEGSIEQLEMIDSLQRLGISYHYKHEIHDILRKIHDQ----    74
                      .* ::*              :. *:: :::* :* ***. *::**::. *  ::

MA_10426551g0010      GIGVGSGDSLIKDLNATALGFRVLRLHQYHVSSGVLENFKDENGQFFCNSTGEQEEENKR    180
AT4G16730             ---HGEIERETQDLHATSLEFILLRQHGFDVSQDAFDVFISETGEFRKTL-------HSD    124
                          *. :   :**:**:* * :** * :.**...:: * .*.*:*  .        :.

MA_10426551g0010      VRSMLSLLRASSISFPGEKVMDEAKAFATDYLNRFLTGRSVT---DVEQSLLREVKYALE    237
AT4G16730             IKGLLSLYEASYFSMDSEFKLKETRIYANKRLSEFVAESSKTICREDETYILEMVKRALE    184
                      ::.:*** .** :*: .*  :.*:: :*.. *..*::  * *   : *  :*. ** ***

MA_10426551g0010      FPWHCSVPRSEARSFIEIYGQNDSWLKSIMNKKILELAKLDFNILQSTHQKDLQLISRWL    297
AT4G16730             TPYHWSIRRLEARWYINVYEKKH-----EMNPLLLEFAAIDFNMLQANHQEELKLISSWW    239
                       *:* *: * *** :*::* ::.      **  :**:* :***:**:.**::*:*** *

MA_10426551g0010      TDSD-IAQLDFYRKRHVELFFWVATGAFEPEFSSSRIAFTKVGTMITFLDDIYDTHGNLD    356
AT4G16730             NSTGLMKQLDFVRDRITESYFWTIGIFYEPEFKYCRKILTKIFMLIVIMDDIYDIYGTLE    299
                      ..:. : **** *.* .* :**.    :****. .*  :**:  :*.::***** :*.*:

MA_10426551g0010      ELKIFTEAVKRWDLSLIDCLPEYMKIALKFFLKTHDELVDEVMRTQGRDMASYIRKNAWE    416
AT4G16730             ELELFTNVVEKWDVNHVERLPNYMRMCFLFLYNEINQIGYDVLRDKGLNVIPYLK-QVWT    358
                      **::**:.*::**:. :: **:**::.: *: :  :::  :*:* :* ::  *:: :.*

MA_10426551g0010      RFIEASMQDAEWRVSGHVPTFDEYIKHAQTTTGLCAVNLITLLLMGPLLPDNILEQIYSP    476
AT4G16730             DLFKTFLTESKWYKTGHKPSFEEYMQNGVISSSVPTILLHLF----SVLSDHISDQTLTD    414
                       :::: : :::*  :** *:*:**:::.  :::: :: *  :     :* *:* :*  :

MA_10426551g0010      SKFQH-L---IELTTRLTDDARDFEAEKEHGELASSVECYLKDNPEFTVEDALNHTTGIV    532
AT4G16730             DSKNHSVVRSCATILRLANDLATSTEEMARGDSPKSVQCYMYETRASEE-EARRHM----    469
                      .. :* :        **::*      *  :*:  .**:**: :.      :* .*

MA_10426551g0010      YRSLKELNWEFIKHDS-------VPLCCKKFVFNFARSMQLFFKDTDGFSNSTKE-IKDQ    584
AT4G16730             -QSMISDSWDIINSDLKTAHTSSLPRGFLAAAANLNRVVQCIYRHGDGHGSPEKTKTVDY    528
                       :*: . .*::*: *        :*      . *: * :* :::. **...  *    *

MA_10426551g0010      IFKVLIDQVPV 595
AT4G16730             IQSVLFNPVPL 539
                      * .**:: **:
```

D MA_10426551_306, cline 0.3
G > C
GAT > CAT
Asp(D) > His(H)
Asp33His
Polar hydrophilic > Polar hydrophilic

W MA_10426551_337, cline 0.5
G > T
TGG > TTG
Trp(W) > Leu(L)
Trp43Leu
Non-polar hydrophobic > Non-polar hydrophobic

**Figure S13** Alignment between Norway spruce BARELY ANY MERISTEM 1 (BAM1) (MA_40935g0020) and *Arabidopsis thaliana* BAM1 (AT5G65700).

```
MA_40935g0020      ------------------------------------------------------------ 0
AT5G65700          MKLFLLLLFLLHISHTFTASRPISEFRALLSLKTSLTGAGDDKNSPLSSWKVSTSFCTWI 60


MA_40935g0020      ------------------------------------------------------------ 0
AT5G65700          GVTCDVSRRHVTSLDLSGLNLSGTLSPDVSHLRLLQNLSLAENLISGPIPPEISSLSGLR 120


MA_40935g0020      ------------------------------------------------------------ 0
AT5G65700          HLNLSNNVFNGSFPDEISSGLVNLRVLDVYNNNLTGDLPVSVTNLTQLRHLHLGGNYFAG 180


MA_40935g0020      ------------------------------------------------------------ 0
AT5G65700          KIPPSYGSWPVIEYLAVSGNELVGKIPPEIGNLTTLRELYIGYYNAFEDGLPPEIGNLSE 240


MA_40935g0020      ------------------------------------------------------------ 0
AT5G65700          LVRFDGANCGLTGEIPPEIGKLQKLDTLFLQVNVFSGPLTWELGTLSSLKSMDLSNNMFT 300


MA_40935g0020      ------------------------------------------------------------ 0
AT5G65700          GEIPASFAELKNLTLLNLFRNKLHGEIPEFIGDLPELEVLQLWENNFTGSIPQKLGENGK 360


MA_40935g0020      ------------------------------------------------------------ 0
AT5G65700          LNLVDLSSNKLTGTLPPNMCSGNKLETLITLGNFLFGSIPDSLGKCESLTRIRMGENFLN 420


MA_40935g0020      ------------------------------------------------------------ 0
AT5G65700          GSIPKGLFGLPKLTQVELQDNYLSGELPVAGGVSVNLGQISLSNNQLSGPLPPAIGNFTG 480


MA_40935g0020      ------------------------------------------------------------ 0
AT5G65700          VQKLLLDGNKFQGPIPSEVGKLQQLSKIDFSHNLFSGRIAPEISRCKLLTFVDLSRNELS 540


MA_40935g0020      ---------------------------------MQSLTSVDFFYNNLSSIVPNGDQFDFF 27
AT5G65700          GEIPNEITAMKILNYLNLSRNHLVGSIPGSISSMQSLTSLDFSYNNLSGLVPGTGQFSYF 600
                                                    ******:** *****.:**. .**.:*

MA_40935g0020      NATPFVGNPGLCGLYLGPCTFHANS-PHTHVMGSSISSTVKMVLVIGLLLCSLVFAFATI 86
AT5G65700          NYTSFLGNPDLCGPYLGPCKDGVAKGGHQSHSKGPLSASMKLLLVLGLLVCSIAFAVVAI 660
                   * * *:***.*** *****.  . .  *     . :*:::*::**:***:**:.**..:*

MA_40935g0020      IKAEALKKSNASISWKLTTFQRLDFTCDDVLECLKEENIIGKGGVGILYKGSMQIGEEFV 146
AT5G65700          IKARSLKKASESRAWRLTAFQRLDFTCDDVLDSLKEDNIIGKGGAGIVYKGVMPNGDLVA 720
                   ***.:***:. * :*:**:************:.***:*******.**:*** *  *: ..

MA_40935g0020      LKRLPGIGRVYYHDHGYVAEIQTMERFGIAIW---------------------------- 178
AT5G65700          VKRLAAMSRGSSHDHGFNAEIQTLGRIRHRHIVRLLGFCSNHETNLLVYEYMPNGSLGEV 780
                   :*** .:.*   ****: *****: *:

MA_40935g0020      ------------------------------------------------------------ 178
AT5G65700          LHGKKGGHLHWDTRYKIALEAAKGLCYLHHDCSPLIVHRDVKSNNILLDSNFEAHVADFG 840


MA_40935g0020      ------------------------------------------------------------ 178
AT5G65700          LAKFLQDSGTSECMSAIAGSYGYIAPEYAYTLKVDEKSDVYSFGVVLLELVTGRKPVGEF 900


MA_40935g0020      ------------------------------------------------------------ 178
AT5G65700          GDGVDIVQWVRKMTDSNKDSVLKVLDPRLSSIPIHEVTHVFYVAMLCVEEQAVERPTMRE 960


MA_40935g0020      --------------------------------------------     178
AT5G65700          VVQILTEIPKLPPSKDQPMTESAPESELSPKSGVQSPPDLLNL     1003
```

Y MA_40935_29209, cline 0.4
A > C
TAT > TCT
Tyr(Y) > Ser(S)
Tyr158Ser
Polar hydrophilic > Polar hydrophilic

G MA_40935_29257, cline 0.4
G > A
GGC > GAC
Gly(G) > Asp(D)
Gly174Asp
Non-polar hydrophobic > Polar hydrophilic

**Figure S14** Alignment between Norway spruce REDUCED VERNALIZATION RESPONSE 1 (VRN1) (MA_10430774g0010) and *Arabidopsis thaliana* VRN1 (AT3G18990).

```
MA_10430774g0010      --------------------RIPPAFVGKLRNQANGNVILEGPSSQRWSVKLCYTSVGME   40
AT3G18990             MPRPFFHKLIFSSTIQEKRLRVPDKFVSKFKDELSVAVALTVPDGHVWRVGLRKADNKIW   60
                                          *:*  **.*:::: .  * *  *..: * * *  :.  :

MA_10430774g0010      FQKGWEKFVCDHRLELGDFLVFKYTSKSYFTVQIFGRSGCEKRNVFN--M----KNFGAS   94
AT3G18990             FQDGWQEFVDRYSIRIGYLLIFRYEGNSAFSVYIFNLSHSEINYHSTGLMDSAHNHFKR-   119
                      **.**::**  : :.:* :*:*:* .:* *:* **. * .* .    .  *    ::*

MA_10430774g0010      LLSLEEKANRDAQDNFPDTLVKSFDADRYQVVPNHTNSQMPVILYTDEDSNDRAMGTFKF   154
AT3G18990             ARLFEDLEDEDAEVIFPSSVYPSPLPE--STVPANKG-------YA-----SSAIQT-LF   164
                         :*:  :.**:  **.::  *   :  ..** :..       *:      . *: *  *

MA_10430774g0010      KQSVSETCPDSAKHPCHGIAARTIKNTESCQPSFAGKKHAIVTSAETGGKSYMQAYYVSC   214
AT3G18990             TGPVKAEEP----TPTPKIPKKRGRKKKNADPEEINSSAP--RDDDPENRSKFYESASAR   218
                      .  *.   *     *   *  :  ::.:..:*.  ...     . :  .:* :     :

MA_10430774g0010      RRAVTEAERNRALKAANSFKSNKPLALIVMKPSHVYQGFWLALPKRLGNCFQLPKGRYQV   274
AT3G18990             KRTVTAEERERAINAAKTFEPTNPFFRVVLRPSYLYRGCIMYLPSGFAEKYLSG--ISGF   276
                      :*:**  **:**::**::*: .:*:  :*::**::*:*  : **. :.: :         .

MA_10430774g0010      TLLGPANKEWVVQYIGDRGRAGLSGGWKHFAIDNNLEEGDVCIFELVNKNKYTFKVYIFR   334
AT3G18990             IKVQLAEKQWPVRCLYKAGRAKFSQGWYEFTLENNLGEGDVCVFELLRTRDFVLKVTAFR   336
                        :  *:*:* *: :  . *** :* ** .*:::*** *****:***:....:.:**  **

MA_10430774g0010      VIKNCIPCKRMEGTKLNEKNGRGRKSIALPSLSLGEKANCGAQNKCPDTLVNSSDADQNQ   394
AT3G18990             VNEYV-------------------------------------------------------   341
                      * :

MA_10430774g0010      IVAKCTNSKLLISLDSDEELDDQEMATHKFEQSVAETSLNHAEPGCCGIAARTVMNTESC   454
AT3G18990             ------------------------------------------------------------   341


MA_10430774g0010      QPSFAGKKCKMVTSAENKGKKCKIVTCAENEGKKHKIITSAENKRKKRKIVTSAENEGKK   514
AT3G18990             ------------------------------------------------------------   341


MA_10430774g0010      RKIVTSAENKGKKGKIVTSAENEGKKGKIVTSTENEGKSYLGIYYVSCRRAVTQAERKKA   574
AT3G18990             ------------------------------------------------------------   341


MA_10430774g0010      LETANSFKSSKPFSLLIMTPSHVYQGFYLSFPRRFEACFQLPRERMEVTLLGPTNKEWQV   634
AT3G18990             ------------------------------------------------------------   341


MA_10430774g0010      QYLGDRSHPGLSGGWRVVKKCTPFKKMEGTKVNDKDVQCRKIIADPVNHFKKMKVSTMST   694
AT3G18990             ------------------------------------------------------------   341


MA_10430774g0010      KKIATNSAVPASGRRIPDILEQGIVPKKEEKQVRKKKDHSSSPNNIIEMDKSSWDSRIDR   754
AT3G18990             ------------------------------------------------------------   341


MA_10430774g0010      RSSPRFNRVCNSSEHPKQLSRPAFTNRKRTIDSDTVLLSAKTNAQLDDTNGEADLIIISD   814
AT3G18990             ------------------------------------------------------------   341


MA_10430774g0010      SNDVAGPSSIIPSQESIHERPSLTKSNVQVVKQENQA 851
AT3G18990             ------------------------------------- 341
```

S MA_10430774_16836, cline 0.4
C > T
TCA > TTA
Ser(S) > Leu(L)
Ser641Leu
Polar hydrophilic > Non-polar hydrophobic

**Figure S15** Alignment between Norway spruce JMJC DOMAIN-CONTAINING PROTEIN 27 (JMJ27) (MA_115863g0010) and *Arabidopsis thaliana* JMJ27 (AT4G00990).

```
MA_115863g0010      MGPLGPSSSHMALGTPCLGPVGPQDIISLVIPRIGDNMSHRGELYTSVPQKEWDVYPEFF      60
AT4G00990           ------------------------------------------------------------      0


MA_115863g0010      DLDLSCAALNIDDHELRNSSSIPRSTIEHAFCSIFVDEDEDEMRWIYAICGNADELACGP      120
AT4G00990           ------------MEKMRGKRIRPRDSG------ELVEDGRSESERKT-RKKENDVVSKGR      41
                                 .::*..   **.:        :*::...* .       : * :: *

MA_115863g0010      LFFFGSESWLTMICFNSSISGKSISVANSEARDVSVKNVESPGEVESKMCHQCQK-NDKG      179
AT4G00990           I---GRG------RGRGEVSKRSIEIDISNP-EKDI-KPDGSRKCLGSTCHHCKILTSES      90
                    :   *          ...:* :**.:  *:  : .: : :.  :  .. **:*:  ..:.

MA_115863g0010      RVVSCYRCKRRRYCIPCLTRWYPGQSEEDIQKACPVCRGNCNCKACLRGDGVMKASTNKT      239
AT4G00990           DLIFCSKCNKKCYCFDCIKRSYSERTHEEVRAACPFCMMTCICRACLRLPLVIKPPSEKD      150
                     :: * :*::: **: *:.* *  ::.*::: ***.*  .* *:****   *:*  ::*

MA_115863g0010      YNADKIRYLHYLLSMVLPMLKQIH------------------------------------      263
AT4G00990           T-DVKLKQLQYLLVKVLPVLKDIYTEQNRELEIESTIRGHPVTEANIKRCKLDPSERIYC      209
                        *:: *:***  ***:**:*:

MA_115863g0010      ---------YEQSCP--NCSYDLCVTCCRELREGFQPGGDEADSAQQCLTERTYFESRDN      312
AT4G00990           DLCRTSIANFHRSCPNKNCSVDICLSCCKELSEGFHQERDGKKNAEGKG-----Y-----      259
                             :.:***  *** *:*::**:** ***:   *  ..*:        :

MA_115863g0010      LPKTLKVPRGRQGWELQAHSEQNNIFGPLHLPPDWKANSNGSIPCPPKEFGGCGSHLLDL      372
AT4G00990           ---ECRIPAG--------QGKDSDAYVPLHF-STWKLNSDSSIPCPPKECGGCGTSTLEL      307
                         ::* *        :.::.: : ***:   ** **:.******** ****:  *:*

MA_115863g0010      KCIFKINWIAKLEKNAEEMASSCKVPEASKVSRPCTSCFKLDFHVKSGLSDSKLRRASYR      432
AT4G00990           RRLWKRDWVEKLITNAEKCTLNFR-PTDVDIVHECSSCS--------TNSDSIRRQAAFR      358
                    : ::* :*: ** .***: : . : *   .: : *:**            ***  *:*::*

MA_115863g0010      EGDSDNFLYCPTEQDIKDGGLEHFQKHWLRGEPVIVQNALQGSSGLSWEPMVMWRAVRET      492
AT4G00990           KNAHDNFLYSPNAVDLAEDDIAHFQFHWMKAEPVIVRNVLEKTSGLSWEPMVMWRACREM      418
                    :.  *****.*.  *: :..: *** **::.*****:*.*: :************* **

MA_115863g0010      NGKF---TADTKTVKAIDCLDWCEVEINIHHFFKGYLEGRMHKNDWPEMLKLKDWPPSNF      549
AT4G00990           DPKRKGTEEETTKVKALDCLDWCEVEINLHQFFEGYLEGRMHKNGWPEMLKLKDWPPSDL      478
                    : *      :*..***:***********:*:**:**********.*************::

MA_115863g0010      FEERLPRHGAEFVSALPFQEYTHPECALLNLASKIPDHCLKPDLGPKTYIAYGSHEELGR      609
AT4G00990           FEKRLPRHNAEFIAALPFFDYTDPKSGILNLATRFPEGSLKPDLGPKTYIAYGFHEELNR      538
                    **:*****.***::****  :**.*:..:****:::*: .************* ****.*

MA_115863g0010      GDSVTKLHCDMSDAVNVLTHTAEVKFPIWQQDKIEKMRESFRSLDLKELFGDSNKVSGTV      669
AT4G00990           GDSVTKLHCDISDAVNVLTHTAKVEIPPVKYQNIKVHQKKYAEAMLQ-----KQQYSGQV      593
                    **********:***********:*::*  : ::*:  ::.: .  *:     .:: ** *

MA_115863g0010      REEDSGKSANKMESEAEASNISG--SDSITNDNSPSPQDT-ELPNEIQSKVAGDTTDYME      726
AT4G00990           KEASELENKSMKEVDESKKDLKDKAANEEQSNNSSRPSGSGEAEKVIISKEDNPTQPAVS      653
                    :* .. :. .   * : . .::..  ::.  .:**  *..: *  : * **  . *   :.

MA_115863g0010      DDLQDTNAVKSDSLRKYEGPSDDWKEIGTMKSKVKENISHKEENTDVTVGGALWDIFRRQ      786
AT4G00990           T------SVESIQEQKLDA--------------PKETDGNTNERSKAVHGGAVWDIFRRE      693
                           :*:* . :* :.              **. .:.:*.:... ***:******:

MA_115863g0010      DAPKLQCYLRKHWKEFRHINGRPVDSIIHPIHDQTFFLNEEHKKKLKEEFSVEPWTFQQH      846
AT4G00990           DVPKLIQFLKRHEHEFRHFNNEPLESVIHPIHDQTMFLSDSQKKQLKEEFDIEPWTFEQH      753
                    *.***  :*::* :****:*..*::*:*******:**.:.:**:*****.:*****:**

MA_115863g0010      LGEAVLIPAGCPHQVRNLKHNQDHSSDEQLGKKQDNVTKHEEEEEFYVCIVMNKDSRSEE      906
AT4G00990           LGEAVFIPAGCPHQVRNRQSCIKVALDFVAP------------ESVEEECLRLTQEFRRLP      801
                    *****:*********** :   . : *                *..  *: :.:: *

MA_115863g0010      DKKKTLREVEDEVVKKKTLKEGEDQVVKRDSASSEKQTQNFDMLDHLDVKEVSLVNMSSF      966
AT4G00990           K----------------------------DHSS---------SEDKLELKKIALYAASSA      824
                    .                            * :*           *:*::*:::*   **

MA_115863g0010      MSMVEGLTIQSTLALIQLVSQCPAKAVFGLRTAVSVIVLGSRRSVEGSRGLGQPRKGSYR      1026
AT4G00990           IREVKGLMQSSRRSDT-------------------------------------------      840
                    :  *:**  .*  :

MA_115863g0010      PVEGPASLGEAAAGSKEGSASLGEAAAGNRRLDLLIEDHCRSCDSWQQRKGLLQPNSQQF      1086
AT4G00990           ------------------------------------------------------------      840
```

```
MA_115863g0010      EAVTASVTAALRQPLRTWKISTVELEKGWKNSATAQLSTSMESKEGFRST   1136
AT4G00990           --------------------------------------------------   840
```

T MA_115863_1213, cline 0.3
A>G
ACT > GCT
Thr(T) > Ala(A)
Thr1103Ala
Polar hydrophilic > Non-polar hydrophobic

**Figure S16** Alignment between Norway spruce LATE MERISTEM IDENTITY2 (LMI2) (MA_128244g0010) and *Arabidopsis thaliana* LMI2 (AT3G61250).

```
MA_128244g0010      MGRAPCCEKTGLKKGPWTSEEDKILVNYIQENGHGSWRALPKLAGLLRCGKSCRLRWTNY      60
AT3G61250           MGRTPCCDKIGLKKGPWTPEEDEVLVAHIKKNGHGSWRTLPKLAGLLRCGKSCRLRWTNY      60
                    ***:***:* ******** ***::** :*::*******:*********************

MA_128244g0010      LKPDIKRGSFTQEEERTIIQLHAILGNRWSVIASQLLGRTDNEIKNFWNTHLKKRLLRMG      120
AT3G61250           LRPDIKRGPFTADEEKLVIQLHAILGNRWAAIAAQLPGRTDNEIKNLWNTHLKKRLLSMG      120
                    *:****** ** :**: :***********:.**:** *********:********** **

MA_128244g0010      IDPSTHAPRTDANPSSPQAFKALNLTRHKAQWENARLEAEVRLSRESLFMSSNSAAGYRT      180
AT3G61250           LDPRTHEPLPSYGLAK--QAPSSPTTRHMAQWESARVEAEARLSRESMLFSPSFYSGVV-      177
                    :** ** *  . . :.     :   *** ****.**:***.******:::* .  :*

MA_128244g0010      NNTVELITMPDDNAPTDYFLKIWSSEAGKAFRKESSCRIINKSCNNSCHVFD--------      232
AT3G61250           ------------KTECDHFLRIWNSEIGEAFRNLAPLDESTITSQSPCSRATSTSSALLK      225
                                ::  *:**:**.** *:***: :     . :.:. *

MA_128244g0010      -LNHQNGGQDDQLLSSDTPKTPISSSLVSSLYCVSTQQCCYRSLNPDDTDKDNDSQHCNS      291
AT3G61250           SSTNSWGGKEVTVAIHG---------------------SDYSPYSN---DLEDDSTD---      258
                      .:. **::  :   .                     . *   .    * ::** .

MA_128244g0010      ASAEWIGDSSVKTPDVLSCFADMTMGQTERKVHKDSSSSEELGPAHVHCTGFSDLVQDFP      351
AT3G61250           --------------SALQLLLDFPISDDDMSFLEENIDSYS----QAPPIGLVSMVSKF-      299
                                  ..*. : *: :.: : .. ::. .* .    :.   *: .:*..*

MA_128244g0010      DNNNSIGCMIPELSTSFRQQSTWPWSELNLEDSKDHWNNVSVTTRASAHG   401
AT3G61250           --------------------------------------------------   299
```

C MA_128244_7736, cline 0.3
T > A
TGT > AGT
Cys(C) > Ser(S)
Cys217Ser
Polar hydrophilic > Polar hydrophilic

**Figure S17** Alignment between Norway spruce LATE MERISTEM IDENTITY2 (LMI2) (MA_118448g0010) and *Arabidopsis thaliana* LMI2 (AT3G61250).

```
MA_118448g0010      MDRAAPCCAKKGLKKGPWSHEEDQILLNYIQRNGHASWRALPELAGLLRCGKSCRLRWTN      60
AT3G61250           -MGRTPCCDKIGLKKGPWTPEEDEVLVAHIKKNGHGSWRTLPKLAGLLRCGKSCRLRWTN      59
                        :*** * *******: ***::*: :*::***.***:**:*****************

MA_118448g0010      YLRPDIKRGSFTEQEERTIIQLHAILGNRWSSIASQLPGRTDNEIKNFWKTRLQKRLLKM      120
AT3G61250           YLRPDIKRGPFTADEEKLVIQLHAILGNRWAAIAAQLPGRTDNEIKNLWNTHLKKRLLSM      119
                    ********* ** :**: :***********::**:***********:*:*:*:****.*

MA_118448g0010      GLDPSTHVPIAEAKPSAPEGYKALAVTRHKAQWESARLEAEARHSSESLIMSSTSTAGYH      180
AT3G61250           GLDPRTHEPLPSYGLA--KQAPSSPTTRHMAQWESARVEAEARLSRESMLFSPSFYSGVV      177
                    **** ** *: .    :  :    :   .*** *******:***** * **:::* :  :*

MA_118448g0010      TVPSDNPPADYFLKIWNSEAGKAFRKENHHPNGGGDDPFLSSEIPLSSVSSLCCVKTPEV      240
AT3G61250           -----KTECDHFLRIWNSEIGEAFRNLAPLDES-----TITSQSPCSRATSTS----SAL      223
                         :  .*:**:***** *:***:      :.       ::*: * * .:* .       :

MA_118448g0010      ISCQPDITMGQTE-DKATEELG---------------FPEPLLDFLDTDDNNNNTNGIQS      284
AT3G61250           LKSSTN-SWGGKEVTVAIHGSDYSPYSNDLEDDSTDSALQLLLDFPISDDDMSFL----E      278
                    :... : : * .*   * .  .                 : ****  :**: .       .

MA_118448g0010      ELRTDFLHQPASPWPELLDEDYWINIAFSQN   315
AT3G61250           ENIDSYSQAPPIGLVSMVS-KF---------   299
                    *   .: : *     .::. .:
```

G MA_118448_6847, cline 0.3
G > A
GGG > AGG
Gly(G) > Arg(R)
Gly140Arg
Non-polar hydrophobic  > Polar hydrophilic

**Figure S18** Alignment of three Norway spruce homologs of LATE MERISTEM IDENTITY2 (LMI2)

```
MA_128244g0010      -MGRAPCCEKTGLKKGPWTSEEDKILVNYIQENGHGSWRALPKLAGLLRCGKSCRLRWTN      59
MA_118448g0010      MDRAAPCCAKKGLKKGPWSHEEDQILLNYIQRNGHASWRALPELAGLLRCGKSCRLRWTN      60
MA_6285g0010        -MGRAPCYEKTGLKKGTWSPEEDKILVNYIENNGHGSWRALPKLAGLLRCGKSCRLRWSN      59
                        ***  *.***** *: ***:**:***:.***.******:***************:*

MA_128244g0010      YLKPDIKRGSFTQEEERTIIQLHAILGNRWSVIASQLLGRTDNEIKNFWNTHLKKRLLRM      119
MA_118448g0010      YLRPDIKRGSFTEQEERTIIQLHAILGNRWSSIASQLPGRTDNEIKNFWKTRLQKRLLKM      120
MA_6285g0010        YLRPDIKRGSFTREEERTIIQLHAILGNRWSIIASQLPRRTDNEIKNFWNTNLKKRLLKM      119
                    **:*********.:**************** *****  **********:*.*:****:*

MA_128244g0010      GIDPSTHAPRTDANPSSPQAFKALNLTRHKAQWENARLEAEVRLSRESLFMSSNSAAGYR      179
MA_118448g0010      GLDPSTHVPIAEAKPSAPEGYKALAVTRHKAQWESARLEAEARHSSESLIMSSTS-----      175
MA_6285g0010        GLDPSTHAPKEEAKPTAPHGYEALALTRHKTQWENARLEAEARLSRESLIMSFDSATPI-      178
                    *:*****.*  :*:*::*..::** :****:***.******.* * ***:**  *

MA_128244g0010      TNNTVELITMPDDNAPTDYFLKIWSSEAGKAFRKESSCRIINKSCNNSCHVFDLNHQNGG      239
MA_118448g0010      ---TAGYHTVPSDNPPADYFLKIWNSEAGKAFRKEN------------------HHPNGG      214
MA_6285g0010        -KSTVEIITIPNDNPPADYFLKLWNSEIGKAFRNEN------------------HHPNGG      219
                       *.   *:*.** *:*****:*.** *****:*.                  :* ***

MA_128244g0010      QDDQLLSSDTPKTPISSSLVSSLYCVSTQQCCYRSLNPDDTDKDNDSQHCNSASAEWIGD      299
MA_118448g0010      GDDPFLSSEIPL-----SSVSSLCCV----------------------------------      235
MA_6285g0010        -NDPVFSSEVPL-----SSVSNMCCASTQHSCDQSLNPNDEDKDNGSQQCNSSSFEWIGD      273
                     :* .:**: *      * **.: *.

MA_128244g0010      SSVKTPDVLSCFADMTMGQTERKVHKDSSSSEELGPAH-VHCTGFSDLVQDFPDNN---N      355
MA_118448g0010      ---KTPEVISCQPDITMGQTEDKAT------EE---------LGFPEPLLDFLDTDDNNN      277
MA_6285g0010        CNVKTPEELSCFPGMALGQTEDKVA------EELGPAQIVHYIGFPELLLDFPDNDDKNN      327
                       ***: :**  .:::**** *.       **           ** : : ** *.:   *

MA_128244g0010      SIGCMIPELSTSFRQQSTWPWSELNLEDSKDHWNNVSVTTRASAHG 401
MA_118448g0010      NTNGIQSELRTDFLHQPASPWPELLDE---DYWINIAFSQN----- 315
MA_6285g0010        NTNSFQSELSTAFQQQSASPWPELHDE---EYWSNMSFIDQASADT 370
                    . . :  ** * * :* : ** **  *   ::* *::.  .
```

**Figure S19** Alignment between Norway spruce FLOWERING CONTROL LOCUS A (FCA) (MA_5232462g0010) and *Arabidopsis thaliana* FCA (AT4G16280).

```
MA_5232462g0010      MQLQYPPLQ--------QH-----------PTHTMAHPQQNQIVTTPWDNKQPQAHLDQP      41
AT4G16280            MRDEYRQSRGCGFVKYSSKETAMAAIDGLNGTYTMRGCNQPLIVR--FA------EPKRP      52
                     *: :*   :        .:            *:**   :*  **   :       . .:*

MA_5232462g0010      KPISMAPTMPPTNTTQPIAAATSVVVDGGPAPT-----------LTCNWTLHTSPEGKTY      90
AT4G16280            KPGESREMAPPVGLG-----SGPRFQASGPRPTSNFGDSSGDVSHTNPWRPATSRNVGPP      107
                     ** .     **..       :   .  .** **            *  *   ** :

MA_5232462g0010      YYNSVTGESQWEKPAELTSFEQQKEQPPSQVLQPQA--QMQQTYSYPQFQFQPQ-QLQQS      147
AT4G16280            SNTGIRGAG--------SDFSP----KPGQATLPSNQGGPLGGYGVPPLNPLPVPGVSSS      155
                       ..: * .        :.*.      *.*.  *.        *. * ::  *   :..*

MA_5232462g0010      QILQPAIATMGQK----------PAELTMFERQQQQQ---------PSQILQPQPQIQQP      188
AT4G16280            ATLQQQNRAAGQHITPLKKPLHSPQGLPLPLRPQTNFPGAQAPLQNPYAYSSQLPTSQLP      215
                       **    : **:          *  * :  * * :          *    .  *  * *

MA_5232462g0010      PSQIIQPQSQIQQTPSQILQPPSQIQQTDSHPQFQPQQQQQSQILQPQSQTQQAPSQILQ      248
AT4G16280            PQQNISRA-TAPQTPLNINLRPTTVS--SATVQFPPRS-------------QQQPLQKMQ      259
                     *.* *.      *** :*   *: :.  .:  ** *:.             ** * * :*

MA_5232462g0010      HPSQIQQTDSQILQPPSQIQRT------DSHPQFQPQQQQQSQILQPQSQIQQTDSHRQF      302
AT4G16280            H-------------PPSELAQLLSQQTQSLQATFQSSQQAISQLQQQVQSMQQPNQNL--      304
                     *             ***:: :       . :  ** .**  **: *  ..:** :.:

MA_5232462g0010      QPQQQQQSQIVEPSVRKQY---TIPAIAPAKVAEPTTVTVTTVADGRAPSHRPLTCDWNE      359
AT4G16280            -PLSQ-----NGRAGKQQWAGSAIPRVASTTGSTPV-SYVQTAAPAVSQSVGSVKCTWTE      357
                      * .*        : ::*:   :** :* :. : *.   * *.* . : *   :.* *.*

MA_5232462g0010      HTSPDGLKYYYNKVTGVSQWEKPAELTAFENQQALRAEP---------------------      398
AT4G16280            HTSPDGFKYYYNGLTGESKWEKPEEMIVFEREQQKQQQHQEKPTIQQSQTQLQPLQQQPQ      417
                     ******:***** :** *:**** *: .**.:*  : :

MA_5232462g0010      ------------------------------------------------------------      398
AT4G16280            QVQQQYQGQQLQQPFYSSLYPTPGASHNTQYPSLPVGQNSQFPMSGIGQNAQDYARTHIP      477


MA_5232462g0010      ----------------------------    398
AT4G16280            VGAASMNDISRTQQSRQSPQELMWKNKA    505
```

T MA_5232462_697, cline 0.1
C > T
ACC > ATC
Thr(T) > Ile(I)
Thr338Ile
Polar hydrophilic > Non-polar hydrophobic

A MA_5232462_670, cline 0.1
C > T
GCA > GTA
Ala(A) > Val(V)
Ala347Val
Non-polar hydrophobic > Non-polar hydrophobic

A MA_5232462_465, cline 0.1
C > T
GCG > GTG
Ala(A) > Val(V)
Ala387Val
Non-polar hydrophobic > Non-polar hydrophobic

Q MA_5232462_452, cline 0.1
G > T
CAG > CAT
Gln(Q) > His(H)
Gln391His
Polar hydrophilic > Polar hydrophilic

Q MA_5232462_451, cline 0.2
C > A
CAA > AAA
Gln(Q) > Lys(K)
Gln392Lys
Polar hydrophilic > Polar hydrophilic

**Figure S20** Alignment between Norway spruce TERMINAL FLOWER 1 (TFL1) (MA_178498g0010) and *Arabidopsis thaliana* TFL1 (AT5G03840).

```
MA_178498g0010      MAFDSLREWSRRKLVDSRVIGSELVESG-TGVIDDFIPSALFCVVYGQKKFFWGDKLSLA       59
AT5G03840           -----------MENMGTRVIEPLIMGRVVGDVLDFFTPTTKMNVSYNKKQVSNGHELFPS       49
                                : :.:***   ::     .*:* * *:: : * *.:*:.  *.:*  :

MA_178498g0010      NLQRKPNSVYDFFDEDLASFVNSGFL---FVHALGCYCGEYLHWMATAMIPSKPHKSVVP       116
AT5G03840           SVSSKPRV--EIHGGDLRSFFTLVMIDPDVPGPSDPFLKEHLHWIVTNIPGTTDAT---F       104
                    .:. **.   ::.. ** **..  ::   .    . :  *:***:.* :  :.  .

MA_178498g0010      FQDLAVYEPPRAI-LNHKIVLTMFKINASQ-PIPRPIQRVFFSTRDFSKDNGLGSPVAAS       174
AT5G03840           GKEVVSYELPRPSIGIHRFVFVLFRQKQRRVIFPNIPSRDHFNTRKFAVEYDLGLPVAAV       164
                     :::. ** **     *::*:.:*: :  :  :*.  .* .*.**.*: : .** ****

MA_178498g0010      FYYIDNENELPRISTS    190
AT5G03840           FFNAQRETAARKR---    177
                    *:  :.*.   :
```

D MA_178498_6893, cline 0.2
T > A
GAT > GAA
Asp(D) > Glu(E)
Asp54Glu
Polar hydrophilic > Polar hydrophilic

**Figure S21** Alignment between Norway spruce LIGHT-HARVESTING COMPLEX B7 (LHCB7) (MA_10435703g0010) and *Arabidopsis thaliana* LHCB7 (AT1G76570)

```
MA_10435703g0010      MVMAAMTHG---LFNPFKGGRIPAVSSTLK------HNPTARIHPCKASWQELAGVLIFS     51
AT1G76570             --MALFQEKLSSLSSSY--SSIHSLPRILVSKPRNRIAVTKSRSICRASWQELAGVLVFS     56
                        ** : .    * . :  . * ::   *          *     *:**********:**

MA_10435703g0010      AIPFTAVKAIANSPLAARLQERMQGTKSLAMKEAPEREAATQRARKESVWYGKDRPRWLG     111
AT1G76570             AIPFTAVKAIANSSIGVSLRRRLEEKKKEAVENSSRFKSKAQEARNDSKWYGKERPRWFG     116
                      ************* :.. *:.*:: .*. *:::: . :: :*.**::* ****:****:*

MA_10435703g0010      MIPYDYPEYLDGETPGDYGFDIAGLSKDPMDFDKNFK-----------------------     148
AT1G76570             PIPYDYPPYLTGELPGDYGFDIAGLGKDRLTFDKYFNFEILHARWAMLAALGALIPEVFD     176
                       ****** ** ** ***********.** : *** *:

MA_10435703g0010      ------------------------------------------------------------     148
AT1G76570             LTGTFHFAEPVWWRVGYSKLQGETLEYLGIPGLHVAGSQGVIVIAICQVLLMVGPEYARY     236


MA_10435703g0010      ------------------------------------------------------------     148
AT1G76570             CGIEALEPLGIYLPGDINYPGGTLFDPLNLSEDPVAFEDLKVKEIKNGRLAMVAWLGFYA     296


MA_10435703g0010      ------------------------------- 148
AT1G76570             QAAFTGKGPVQNLVDHVSDPLHNNLIAMLQT 327
```

T MA_10435703_10305, cline 0.4
A > G
ACA > GCA
Thr (T) > Ala (A)
Thr125Ala
Polar hydrophilic > Non-polar hydrophobic

**Figure S22** Gene Ontology (GO) enrichment analysis of top 20 pathways for Biological Process with a p-value cut-off of 0.05 (FDR).
GO enrichment analysis for 2084 Norway spruce genes containing 2936 missense cSNPs, was performed by using ShinyGO 0.85.1, using the Arabidopsis database.

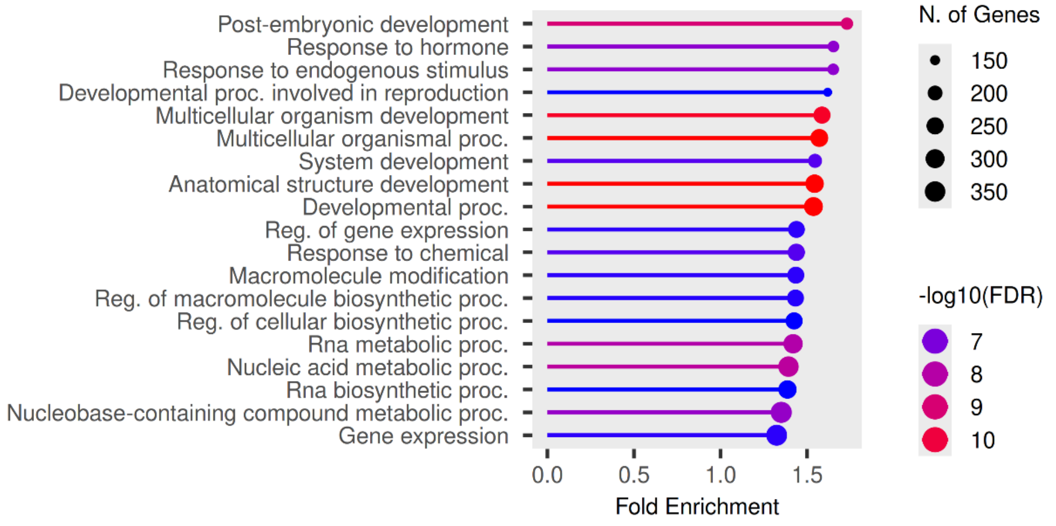


**Figure S23** Gene Ontology (GO) enrichment analysis of top 20 pathways for Molecular Function with a p-value cut-off of 0.05 (FDR).
GO enrichment analysis for 2084 Norway spruce genes containing 2936 missense cSNPs, was performed by using ShinyGO 0.85.1, using the Arabidopsis database.

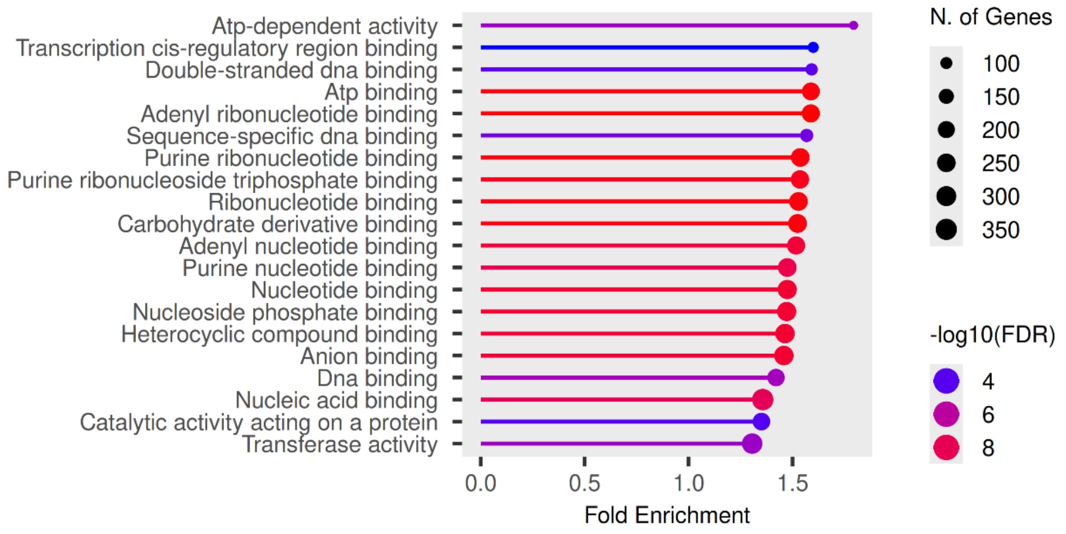


**Figure S24** Gene Ontology (GO) enrichment analysis of top 20 pathways for Cellular Compartment with a p-value cut-off of 0.05 (FDR).
GO enrichment analysis for 2084 Norway spruce genes containing 2936 missense cSNPs, was performed by using ShinyGO 0.85.1, using the Arabidopsis database.

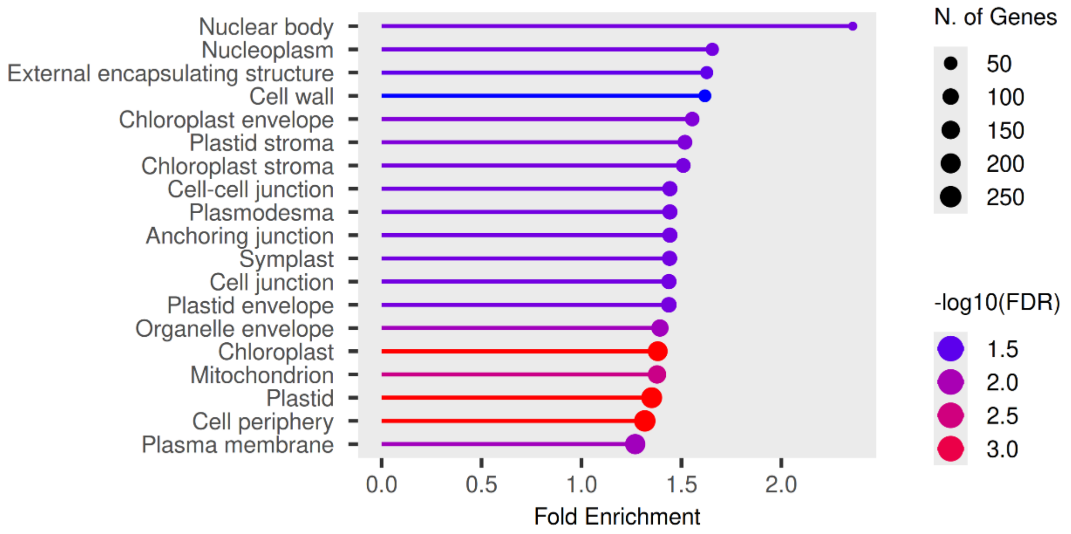

**Figure S25**

**(a) Nucleotide sequence of partial *SQUAMOSA BINDING PROTEIN-LIKE1 (SPL1)* transcript from Norway spruce: Region highlighted in blue is the miR529/miR156-binding site. Nucleotides highlighted in pink represent positions with SNP with very low trend cline and nucleotides highlighted in red represents position with SNP with standard cline**

>**MA_22749g0010** [927 nucleotides]
ATGTTAGGGTCAGTGGATAACAGGCATACTGGCCTTTTGCTAGATGCATCATCCTTTATGCACCCAAGGATTTTTTCAAGTTCAGTTTTGGACGATCAC
GGTGACTTCAAACTTGGACAGGGGAAGGGGACCTGGCCAAAGATTGTTAAATCAGAGGATCCATCAATGTTTGATGGCCATCCACAAACATCAGGTGTT
AATCGGCCACCGTTTACCAATGCTCTGTCTCGTCAAAGTGCTGACAGGTTACTCCAGCTCTTGCAGAGCTCCAAGGCTTCCTCTGTCAATGTTCTCAGT
CAAGGCATTCAACATTATACACACAGCCCAGCAAACCCTCTCACCGAGGCCTTAAGTCTTTCATTTTCACCAGGAACAGGGGTTATTGCTGGTCTAGAA
GTAGCATCGGCTACACAGAATTTATTAGGAGTCTCAGACTCCGGTTGTGCTCTCTCTCTTCTGTCATCGCAGTCATTGGGCTCAAGGGCTCTAGGATCA
GCTTCATTAGATCTGGCCACTCGACCAGGTGCCACATTGGAACAGTTTATGCACGAGAATCATTCTACCATGGATCAAATTTCACTGCAAGGAATGCAA
CATAATTTTAGTCTAAATAGTAGCTCCCAGTCCTCGACAGGTCTCATCTCTAATGGATATCCTCCTCCTGGGGTGAATTTTATAGATAAGGTTCAAGTT
GGGTCACTAACTCCTAATATTGGGGGTATTGGAAACTTTGAAACCCCTATGCATGGCTTTCTGCAAGGACAGGAATTCAGAAGTTCTCAAGATGCAAGA
TCAAAAGCTGTTCGACGTACCATCGATTTAATGCACAGGTCCTCACAGACACAGGGGAATGAATCACAGGGACAATCTGGGATACTGCAGCATGGTAAT
AATGGTTCAATGAACAATTTATGGAATTTCTGTTGA

**(b) Allele frequencies of one of the alleles from SNPs represented in partial nucleotide sequence of *SPL1* across six Norway spruce populations in Sweden**

| CHROM_POS | Ref_To_Alt | Amino acid change | S1_Allele1 | S2_Allele1 | S3_Allele1 | S4_Allele1 | S5_Allele1 | S6_Allele1 | Cline steepness |
|---|---|---|---|---|---|---|---|---|---|
| MA_22749_19139 | 723T>G | Missense ATT>ATG Ile(I)>Met(M) | 0.97 | 0.96 | 0.97 | 0.91 | 0.9 | 0.94 | 0.03 |
| MA_22749_19144 | 728A>G | Missense AAC>AGC Asn(N)>Ser(S) | 0.97 | 0.99 | 0.97 | 0.97 | 0.96 | 0.93 | 0.04 |
| MA_22749_19154 | 738C>T | Synonymous ACC>ACT Thr(T)>Thr(T) | 0.98 | 0.97 | 0.98 | 0.99 | 0.99 | 0.99 | 0.01 |
| MA_22749_19157 | 741T>C | Synonymous CCT>CCC Pro(P)>Pro(P) | 0.06 | 0.05 | 0.11 | 0.06 | 0.1 | 0.09 | 0.03 |
| MA_22749_19187 | 771C>T | Synonymous TTC>TTT Phe(F)>Phe(F) | 0.9 | 0.9 | 0.81 | 0.75 | 0.75 | 0.63 | 0.27 |

**Figure S26**

**(a)** Alignment between Norway spruce STYLISH 1 (STY1) (MA_53545g0010) and *Arabidopsis thaliana* LAF3 (AT3G51060).

```
MA_53545g0010      MEEVRGCRGSELWLIVATVAPAVASTGGASGGPRRTEGGIPEWVSSSSSSSLQLQPRGVI 60
AT3G51060          ------------------------------------------------------------ 0


MA_53545g0010      KERDFSLGFNGGGGGAGGVTHSAGSTSTSHTHHHSRYTLVRPLSYGYPSGIEMAVGGMLG 120
AT3G51060          ------------------------------------------------------------ 0


MA_53545g0010      AGVGESPAGGSGPYAGGPARLGSISGGSGGINIVNRESLRDVVTRAGGGGGGGTEDFVSL 180
AT3G51060          --------------------------------------MAGFFSLDGGGGGGGGGGNNQE 22
                                                         : ...:  *******  .  .

MA_53545g0010      YTRVRREDDPTSAPRGGGSFVAYATTPRTVDDPDHHANIQSSMPVLSSAHHREDDRGVDQ 240
AT3G51060          D--HRSNTN--PPPPVSEAWLWYRN-PNVNAN--ANTNVNANAP---------SSSNAAL 66
                       * : :    *  . ::: * . *..  :   ::*:::. *         .. ..

MA_53545g0010      GEIQVWQQARDLRGDGNQESKEKAMASQMRVMEISGG-QQAPERRSNCELGATGDQSQLS 299
AT3G51060          GTLELWQNHNQQEIMFQHQQHQQ----RLDLYSSAAGLGVGPSNHNQF---------DIS 113
                   * :::**: .: .   :::.:::    :: : . :.*   .*..:.:          ::*

MA_53545g0010      NVENKNGGGGLAVTVAAMPKTNTNPNTITNANSTTCQDCGNQAKKDCRHLRCRTCCKSRG 359
AT3G51060          GETST-AGAGRAAAMMMI----RSGGSGGGSGGVSCQDCGNQAKKDCSHMRCRTCCKSRG 168
                   .  .. .*.* *.::  :     .  .:  .:...:************ *:**********

MA_53545g0010      FDCSTHLKSTWVPAAKRREKQQAEAAAIA-----AGHPRPKTKRARFVSSSNATTTSHTS 414
AT3G51060          FECSTHVRSTWVPAAKRRERQQQLATVQPQTQLPRGESVPKRHREN-------------- 214
                   *:****::***********:**  *:.        *.  ** :* .

MA_53545g0010      TSATTPPRSSDINSTQIAHQDLNFKANLPAEIKAQAVFKCVRVTAIEDGVDEFAYQATVK 474
AT3G51060          ----LPATSSSLVCTRIPSHSGLEVGNFPAEVSSSAVFRCVRVSSVEDGEEEFAYQTAVS 270
                        *  **.: .*:*  :.    .*:***:.:.***:****:::*** :*****::*.

MA_53545g0010      IGGHIFRGVLYDDQRADTSNLHSNLPD---------------LHLGGTTTMASTSAMIDP 519
AT3G51060          IGGHIFKGILYDLGPGSSGGGGYNVVAAGESSSGGGGAQQLNLITAGSVTVATASS-STP 329
                   ******:*:***   ..:..   *:                 *  .*:.*:*::*:   *

MA_53545g0010      TAGGGVYGASSS----NILGGMDYRNPISTQHQTMFDMQCLS----  557
AT3G51060          NLGG--IGSSSAAAATYIDPAALYPTPINTF---MAGTQFFPNPRS  370
                   . **   *:**:     *  .  * .**.*    * . * :
```

G MA_53545g0010, cline 0.6
T > C
GGT > GGC
Gly(G) > Gly(G)
Gly31Gly
Non-polar hydrophobic > Non-polar hydrophobic

**(b) Allele frequencies of one of the alleles from synonymous SNP from *STYLISH 1* (*STY1*) across six Norway spruce populations in Sweden**

| CHROM_POS | Ref_To_Alt | Amino acid change | S1_Allele1 | S2_Allele1 | S3_Allele1 | S4_Allele1 | S5_Allele1 | S6_Allele1 | Cline steepness |
|---|---|---|---|---|---|---|---|---|---|
| MA_22749_19139 | T>C | Synonymous GGT>GGC Gly(G)>Gly(G) | 0.28 | 0.33 | 0.52 | 0.80 | 0.86 | 0.91 | 0.6 |
